# Combinatorial Drug Discovery in Nanoliter Droplets

**DOI:** 10.1101/210492

**Authors:** Anthony Kulesa, Jared Kehe, Juan Hurtado, Prianca Tawde, Paul C. Blainey

## Abstract

Combinatorial drug treatment strategies perturb biological networks synergistically to achieve therapeutic effects and represent major opportunities to develop advanced treatments across a variety of human disease areas. However, the discovery of new combinatorial treatments is challenged by the sheer scale of combinatorial chemical space. Here we report a high-throughput system for nanoliter-scale phenotypic screening that stabilizes a chemical library in nanoliter droplet emulsions and automates the formulation of chemical combinations *en mass* using parallel droplet processing. We apply this system to predict synergy between more than 4,000 investigational and approved drugs and a panel of 10 antibiotics against *E. coli*, a model Gram-negative pathogen. We found a range of drugs not previously indicated for infectious disease that synergize with antibiotics. Our validated hits include drugs that synergize with the antibiotics vancomycin, erythromycin, and novobiocin, which are used against Gram-positive bacteria but are not effective by themselves to resolve Gram-negative infections.

## Main Text

Much of modern drug discovery acts to modulate a specific drug target using a single agent with maximally selective effects, arising from the idea of Paul Ehrlich’s “magic bullet” *(1)*. However, the prevalence of redundancy, feedback, and multifunctionality in biological networks challenges this approach *(2–4)*. Therapeutic strategies comprising multiple drugs in combination have been proposed to exploit network-driven interactions to achieve the desired functional perturbation, reduce toxicity, and prevent or overcome drug resistance *(2–6)*. In particular, combination antimicrobial treatments that overcome drug resistance by targeting known resistance elements (*e.g.* beta-lactamase enzymes) in addition to essential targets make up a substantial fraction of antibiotic treatments in clinical development today *(7)*.

Despite the applicability of novel drug combinations, their identification by high-throughput screening has been slowed by the high complexity, cost, and compound consumption of conventional screening methods *(8)*. For example, testing all pairs of drugs from a modest library of 2,000 drugs (*e.g.* FDA approved drugs) requires almost 2 million pairwise combinations, and far more if compounds are titrated. Experiments of this scale are restricted to specialized labs and facilities that can accommodate the large costs and complexity (total liquid handling steps, logistics of plate layout and workflow design). Additionally, since these screens test each compound across thousands of others, thousands-fold more compound is required than single-compound screening, which can deplete an entire inventory in a single screening experiment. Current methods for combinatorial discovery work around these issues, either by leveraging computational predictions of drug synergies to reduce screening scale, or by combining multiple tests in pools with subsequent deconvolution *(9–11)*.

Here we introduce a strategy for combinatorial drug screening based on droplet microfluidics that unlocks order-of-magnitude improvements in logistical complexity and compound consumption, and reduces demand for capital equipment (**Fig. 1**). Recent advances in droplet microfluidics are making major impacts across the life sciences by processing cells and nucleic acid molecules in high speed serial streams of water-in-oil emulsion droplets but have not yet been fully translated to chemical screening *(12–14)*. Our platform leverages the throughput potential of microfluidic and microarray systems *(15–17)*, and substitutes deterministic liquid handling operations needed to construct combination of pairs of compounds with parallel merging of random pairs of droplets in a microwell device (**Fig. 1**). Unique advantages of this method are that it can be hand-operated at high throughput to eliminate the need for robotic liquid handling, and that assay miniaturization in microwells enables parsimonious use of compound to reduce consumption.

**Fig 1.**
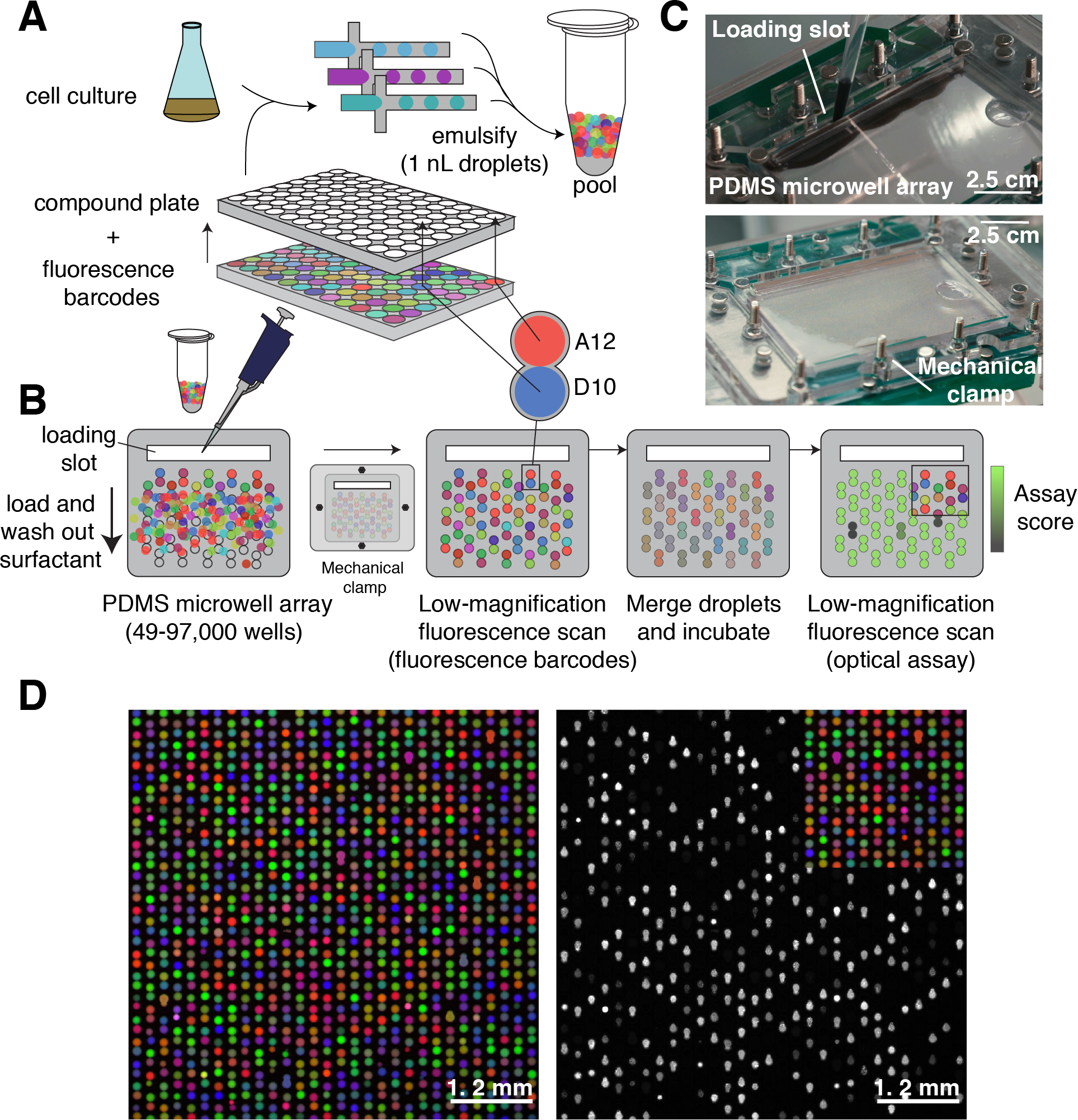
Droplets platform for combinatorial drug screening. (**A**) Compounds, cells, and fluorescence barcodes are emulsified into nanoliter droplets and subsequently pooled. (**B**) A microwell array (**Fig. S1**) pairs random combinations of droplets (**Movie S1**, **S2**). Once loaded, free surfactant is depleted by washing the microwell array to limit compound exchange. Low-magnification epifluorescence microscopy identifies the compounds carried by each droplet. Pairs of droplets in each microwell are merged and incubated (**Movie S3**). A second optical scan reads out a phenotypic assay (*e.g.* cell growth inhibition determined by GFP fluorescence). (**C**) Photographs of microwell array assembly during and after loading (**Fig. S1**, **Movie S1**, **S2**). Scale bars are approximate due to perspective effect. (**D**) Three-color fluorescence micrograph of droplets in microwell array paired with a subsequent assay of growth inhibition of *E. coli* cells, monitored by fluorescence from constitutively expressed GFP. Only 50% of droplet inputs contained cells, therefore a fraction of microwells do not contain cells and as a result do not show GFP fluorescence.

Our platform constructs and assays all pairwise combinations of a set of input compounds (**Fig. 1**). First, the concentrated compounds in well plates are combined with fluorescence barcodes (unique ratios of three fluorescent dyes), cells, and media (**Fig. 1a**). We then emulsify a sample from each well into 20,000 1-nanoliter aqueous droplets in a fluorocarbon oil continuous phase with a stabilizing fluorosurfactant. We use standard multi-channel micropipettes to combine the droplets into one pool, and load the pooled droplets into a microwell array such that each microwell captures two droplets at random (**Fig. 1a-c**, **Fig. S1**, **Movies S1**, **S2**). We then seal the microwell array to the glass substrate to limit microwell cross-contamination and evaporation and fix the assembly by mechanical clamping (**Fig. 1b**, **Fig. S1**). We identify the contents of each droplet by reading the fluorescence of the encoding dyes (95-99% accuracy; **materials and methods**, **Fig. S2**) by low-magnification epifluorescence microscopy (2-4X, 6.5 μm/pixel optical resolution) (**Fig. 1b**, **d**) *(18)*. We then merge all pairs of droplets by applying a high-voltage AC electric field (**Movie S3**), and incubate the microwell array to allow cells to respond to the pair of compounds (**Fig. 1b**) *(19)*. Last, we image the microwell array to read out the assay result (*e.g.* cell growth inhibition) and map this measurement to the pair of compounds previously identified in each well (**Fig. 1b**, **d**).

We designed our platform to overcome a critical challenge in droplet microfluidics that has heretofore prevented cell-based compound screening with hydrophobic small molecules: the exchange of compounds between droplets on assay-relevant timescales *(20–22)*. The dynamic equilibrium of surfactant molecules between the aqueous-oil droplet interface and reverse-micelles in the oil phase can carry small molecules between droplets (**supplementary text**) *(20)*. Our microwell array design limits compound exchange after loading by (i) depleting free surfactant by an oil wash, and (ii) limiting reverse-micelle diffusion between microwells by mechanically sealing the microwell array to a substrate (**Fig. 1a**, **b**, **Fig. S1**). To measure compound cross-contamination on our platform, we monitored the transport of a fluorescent dye (resorufin) from “source” droplets (resorufin) to “sink” droplets (fluorescein, minimal exchange on assay timescales) (**Fig. 2a**) *(20, 21)*. We found that compartmentalization alone (without depletion of free surfactant) limited resorufin transport compared to exchange between pairs of droplets in the same microwell (**Fig. 2b**, **c**). Depleting free surfactant by washing the loaded microwell array prior to sealing further decreased exchange to levels below our detection limit (**Fig. 2d**). While cross-contamination cannot be eliminated in the brief droplet pooling phase prior to loading the microwell array (**Fig. 1a**, **2a**), we expect <5% exchange under screening conditions for compounds no more hydrophobic than resorufin (**supplementary text**, **Fig. 2d**, **Fig. S3**).

**Fig 2.**
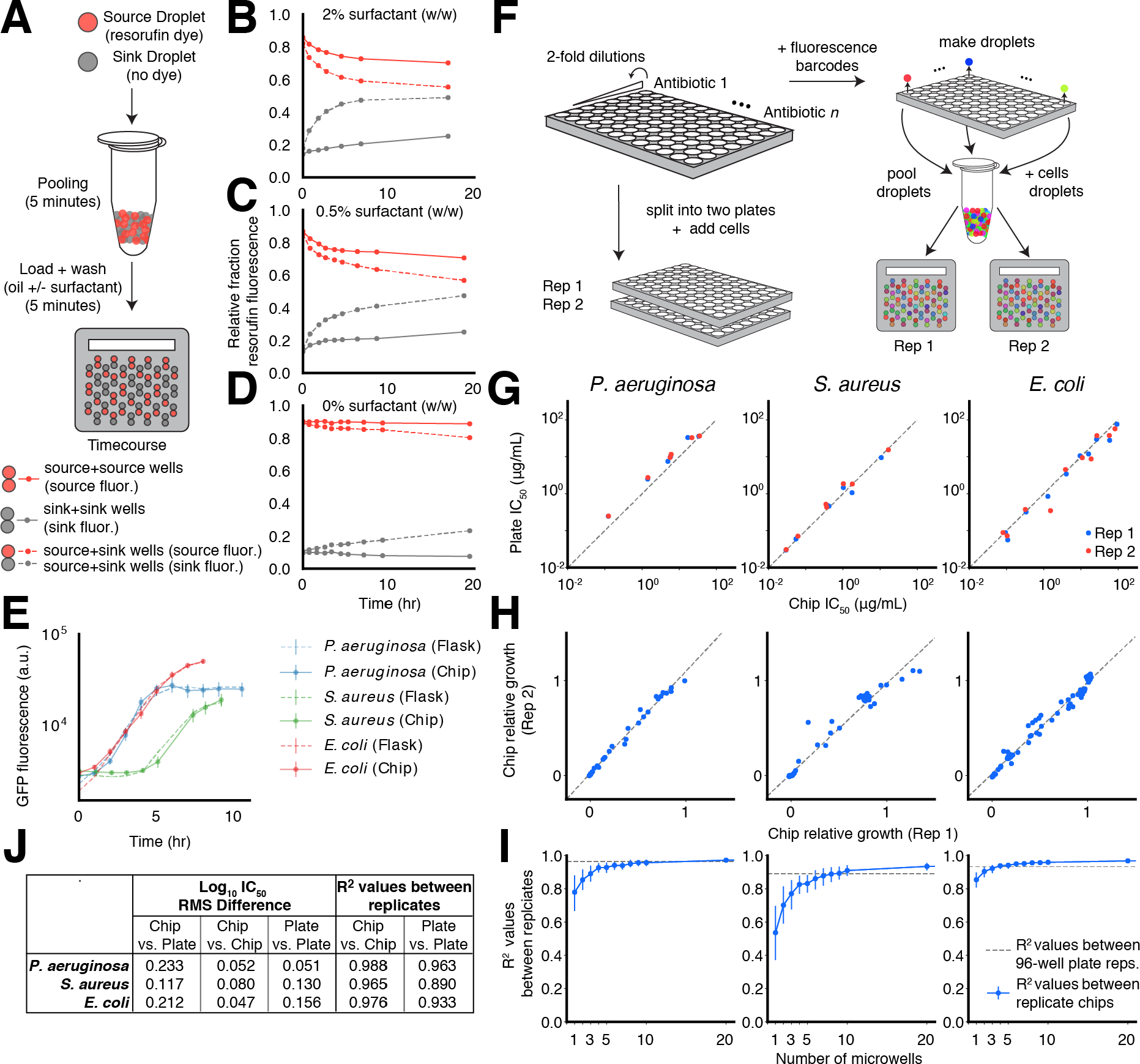
Characterization of droplet platform performance. (**A**) As a model for cross-contamination of screening compounds, we tracked the transfer of the fluorescent dye resorufin (carried by “source droplets”) to empty droplets (“sink droplets”). (**B**, **C**, **D**) Exchange of dye is measured over time by fraction of observed dye fluorescence measured in source droplets (red lines) and sink droplets (gray lines) as a function of surfactant wash concentration (0%, 0.5% and 2% w/w surfactant). The effect of compartmentalization is measured by comparing the rates of dye accumulation in sink droplets when co-compartmentalized in microwells with source droplets (dotted lines) or other sink droplets (solid lines). Transfer that occurred between droplets prior to loading was measured by the fraction of total fluorescence in sink droplets at the first timepoint in the assay (**supplementary text**, **Fig. S3f**). (**E**) We measured cell growth by monitoring accumulation of constitutively expressed GFP in both conventional Erlenmeyer flasks (dotted lines) and the droplet platform (solid lines). Error bars represent standard deviations of microwell measurements. Measurements from Erlenmeyer flask cultures are linearly transformed to the same scale as microwell measurements (**materials and methods**). (**F**) Experimental setup for comparing antibiotic response curves and measuring technical noise in 96-well plate broth culture format and the droplet platform. (**G**) Estimated IC_50_ for each antibiotic compared between 96-well plate and droplet platform formats (**materials and methods**, **Fig. S4**, **S5**, **S6**). Dotted lines show the diagonal. (**H**) Comparison of growth values determined from two technical replicates on the droplet platform (**Fig. S4**, **S5**, **S6**). Dotted lines show line of best fit. (**I**) Relationship between microwell-level replication and technical noise, estimated by bootstrap resampling of the data set in part H. Error bars represent 10-90^th^ percentile bootstrapped R^2^ values. Dotted lines represent R^2^ values between technical replicates in 96-well plate broth culture (**Fig. S7**). (**J**) For data shown in part G, we report root mean square (RMS) differences between log_10_ IC_50_ values for antibiotic growth curves measured in the droplet platform and 96-well plate broth culture format. RMS differences between technical replicates in each format are shown for comparison (**Fig. S7**). For data shown in part H, R^2^ values measured between technical replicates are shown.

As a first application of our platform, we developed fluorescence-based growth inhibition phenotypic screening assays for three model bacterial pathogens often used in antibiotic discovery, *Pseudomonas aeruginosa, Staphylococcus aureus*, and *Escherichia coli*. For each organism, we compared growth dynamics (**Fig. 2e**), antibiotic drug responses (**Fig. 2f**, **g**, **j**, **Fig. S4**, **S5**, **S6)**, and reproducibility of the droplet platform with conventional Erlenmeyer flask and 96-well plate broth culture methods (**Fig. 2f**, **h-j**, **Fig. S7**). Growth dynamics (monitored by constitutive GFP fluorescence) between Erlenmeyer flasks and the droplet platform showed close correspondence, indicating no detectable toxicity or gross physiological impact on the bacteria (**Fig. 2e**). We chose 6-12 antibiotics representing different chemical classes and mechanisms of action, and compared IC_50_ values estimated from five-point dose response curves measured with the droplet platform and the same fluorescence assay in a 96-well plate broth culture format (**materials and methods**, **Fig. S4**, **S5**, **S6**). Overall, culture plates and the droplet platform indicated similar potency for each antibiotic and comparable levels of assay noise (R^2^ values between technical replicates) (**Fig. 2g**-**j**, **Fig. S4**, **S5**, **S6**, **S7**).

High-throughput screening is extremely sensitive to assay noise as hits must be enriched compared to false positives. In the droplet platform, droplets carrying different compounds are paired stochastically in microwells, and noise levels are mitigated by making multiple measurements of the same compound pair across replicate microwells. The number of replicate microwells is a random variable with an expected value determined by the number of possible unique input droplet pair combinations and the number of microwells on a given chip (**materials and methods**, **Fig. S1**), and assay noise can be reduced by choosing a higher replication level at the cost of lower throughput. To explore this relationship, we down-sampled the number of replicate microwells observed per antibiotic dose and compared measurements from two technical replicate microwell arrays. We observed diminishing improvements at microwell replication levels past 5-10 microwells, and this is achieved for 64 unique droplet inputs per microwell array (**materials and methods**, **Fig. 2i**).

To evaluate our ability to detect synergy between compound pairs, we tested a canonically synergistic pair, ampicillin (a beta-lactam antibiotic) and sulbactam (a beta-lactamase inhibitor), against *P. aeruginosa* (**Fig. S8**). Synergy is commonly assessed by crossing all pairs of a dilution series of two drugs in a “checkerboard” assay matrix and quantified via Bliss Independence or the Fractional Inhibitory Concentration (FIC) method (**materials and methods**) *(23, 24)*. Synergy between ampicillin and sulbactam (defined as FIC≤0.5) was detected in both 96-well plate broth culture (FIC≤0.5) and the droplet platform (FIC≤0.25) (**Fig. S8**).

We next applied our system to identify compounds that can potentiate the activity of antibiotic drugs. In the face of rising antibiotic resistance, efforts to develop new classes of antibiotics have yielded little success *(7, 25, 26)*. Unfortunately, many antibiotics such as vancomycin, erythromycin, and novobiocin cannot be used to treat important clinically important Gram-negative pathogens such as *E. coli, P. aeruginosa, A. baumannii,* and *K. pneumoniae* due the impermeability of their outer membranes and numerous efflux systems *(7, 26)*. Previous work suggests that identifying compounds that sensitize drug-resistant pathogens is a promising strategy to broaden the usage of these antibiotics *(27–29)*.

We screened for potentiation of a panel of ten antibiotics with diverse mechanisms and biochemical target localization (each antibiotic titrated across a three-point response curve; **Table S1**) by a “drug repurposing” library of 4,160 compounds against *E. coli* (**Fig. 3a**) *(28, 30)*. This curated repurposing library is composed of tool compounds, compounds with extensive preclinical investigation, and launched drugs *(30)*. We reasoned that hit compounds from screening this library would already have extensive characterization that would expedite potential translation for use against Gram-negative pathogens *(30, 31)*.

**Fig. 3.**
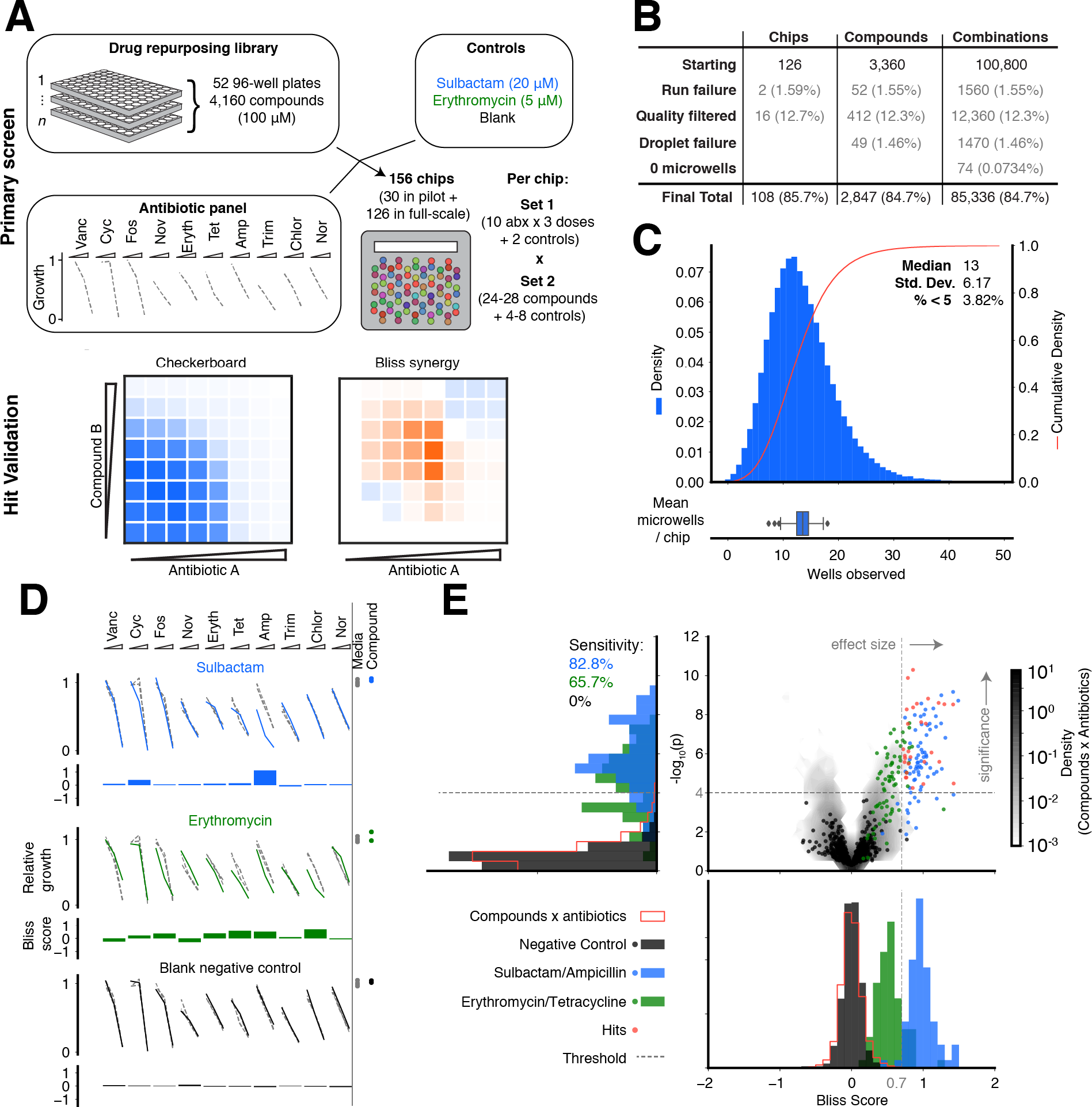
Drug repurposing antibiotic potentiation screen. (**A**) To measure antibiotic potentiation, we generated three-point dose response curves for ten different antibiotics in combination (**Table S1**) with 4,160 compounds (each at single concentration, 100 μM) from a drug repurposing library, as well as positive controls (sulbactam and erythromycin) and negative controls (blank media). Each chip formulated all pairwise combinations of two input sets: i) 3 antibiotics × 10 concentrations (**Table S1**) + 2 controls (32 total); ii) 24-28 compounds + 4-8 controls (32 total). Hits were validated by performing eight-point checkerboard assays to determine Bliss synergy and FIC. (**B)** The final numbers of analyzed combinations in the full-scale screening phase, after accounting for losses and quality filtering (**materials and methods**, **Fig. S9**). (**C**) The histogram (blue bars) and cumulative distribution (red line) of the number of microwells observed for each compound × antibiotic combination. (Bottom panel) The distribution of mean numbers of microwells of all compound × antibiotic combinations on each chip in full-scale phase of screen (Tukey box plot). (**D**) Primary screening data yielded antibiotic response curves in combination with each compound that we compared to response curves of the antibiotic alone (gray, dotted). Growth in the presence of compound alone is shown by the colored dots at right (“Compound”). Growth in the absence of antibiotic and compound is shown by gray dots at right (“Media”). Data from controls (blue: sulbactam; green: erythromycin; black: blank) (**Table S2**) are shown as examples. This comparison is made quantitative by calculating a Bliss Score (**materials and methods**). (**E**) (Top right panel) 28 compound × antibiotic combination hits were determined by thresholding Bliss Scores (gray, shaded contours) for each pair on effect size (Bliss Score > 0.7, gray dotted line) and statistical significance (p < 10^-4^, gray dotted line) (**materials and methods**, **Table S3**, **S4**). (Top left panel) Projection of vertical axis. Sensitivity to positive controls (blue: sulbactam, green: erythromycin) determined by statistical significance threshold (**Table S2**). (Bottom right panel) Projection of horizontal axis. Bliss Score distributions of positive and negative (black: blank) controls (**Table S2**). Histograms are set to 50% opacity to show overlap.

This screening effort resulted in the construction of 4+ million total microwell assays across 156 microwell array chips, and was completed over 10.3 days in two phases (Pilot phase: 800 compounds, 30 chips, 3.33 days; Full-scale phase: 3,360 compounds, 126 chips, 7 days). With a total of 64 unique inputs per microwell array chip (set 1: 10 antibiotics at 3 dose points + 2 controls; set 2: 24-28 compounds + 4-8 controls), each chip run constructed 720-840 combinations of compound × antibiotic (24-28 compound × 10 antibiotics × 3 dose points), 276-378 compound × compound (1/2 × 24 × 23; 1/2 × 28 × 27), 120-240 control × antibiotic combinations (4-8 controls × 10 antibiotics × 3 dose points), and 48-56 compound × control combinations (24-28 compounds × 2 controls) (**Fig. 3a**; **materials and methods**). Pairwise combinations of antibiotics and controls were also constructed.

Our analysis focused on determining compound × antibiotic synergies by evaluating a shift of a three-point antibiotic dose response with and without compound (**Table S1**), quantified by the Bliss synergy metric for each compound-antibiotic pair (**Fig. 3a**, **d**, **materials and methods**). Hits from our screen were then validated in eight-point checkerboard assays and quantified by Bliss Independence and the FIC method (**Fig. 3a**, **materials and methods**).

We evaluated screening performance from our full-scale phase, comprised of 126 microwell array chip runs and 100,800 compound × antibiotic assay points from 3,360 compounds. Dropout can occur due to chip-run failures, failures to produce, load, and classify droplets, or failure to observe any microwells containing a particular compound × antibiotic combination. Of the 126 chip runs, we had two logistical failures and removed 16 runs due to failed controls to yield a final chip passing rate of 85.7% (108 runs) (**Fig. 3b**, **Fig. S9**, **materials and methods**). Droplet production, pooling/loading, or fluorescence barcode assignment failed for 49 compounds (**Fig. 3b**). Overall, of the starting 100,800 compound × antibiotic combinations, 84.7% were successfully measured with an overall median value of 13 replicate microwells (**Fig. 3b**, **c**).

To assess data quality, each chip run was performed with a set of positive (sulbactam × ampicillin; erythromycin × tetracycline) and negative (blank media × all antibiotics) controls to determine the expected sensitivity and false positive rate of the screen (**Fig. 3a**, **d**, **Table S2**). The Bliss Score distribution of all blank media × antibiotic pairs was well-described by a T-distribution, which we used as a null model to calculate p-values for each compound × antibiotic pair (**Fig. S10**, **materials and methods**). To measure sensitivity, each run included one or both positive controls: sulbactam × ampicillin (large effect size, expected Bliss Score ~1) and erythromycin × tetracycline (small effect size, expected Bliss Score ~0.5) (**Fig. 3d**). At an expected false positive rate of 10^-4^ (p-value threshold), we recovered 82.8% of sulbactam × ampicillin controls (n=58/70) and 65.7% of erythromycin × tetracycline controls (n=46/70) (**Fig. 3e**). To call hits from all the compound × antibiotic pairs (**Table S3**), we chose a Bliss Score effect size threshold that separated sulbactam × ampicillin controls from erythromycin × tetracycline controls (Bliss Score > 0.7) (**Fig. 3e**). Using these thresholds to score all pairs yielded 28 hit compound × antibiotic pairs (0.098% of total 28,470) from 20 distinct compounds (0.70% of total 2,847) (**Fig. 3e**, **Table S4**). While we focused analysis on compound × antibiotic pairs, we did identify that one hit compound, pasireotide, also synergized with tedizolid, another compound in the repurposing library run on the same microwell array chip (**Fig. S11**).

We selected 17 hit compound × antibiotic pairs from 11 distinct compounds for confirmation in eight-point checkerboard assays measured in 96-well plate broth culture format (**Fig. 4a**, **Fig. S12**). For comparison, we measured an additional 29 pairs that did not pass Bliss Score and p-value thresholds in the primary screen. Of the hit combinations, 15/17 scored as synergistic by Bliss Independence (88.2%, p = 5.8 × 10^-4^; binomial distribution with 22/46 total tested pairs scoring positive for synergy) (**Fig. 4a**, **Fig. S12**).

**Fig 4.**
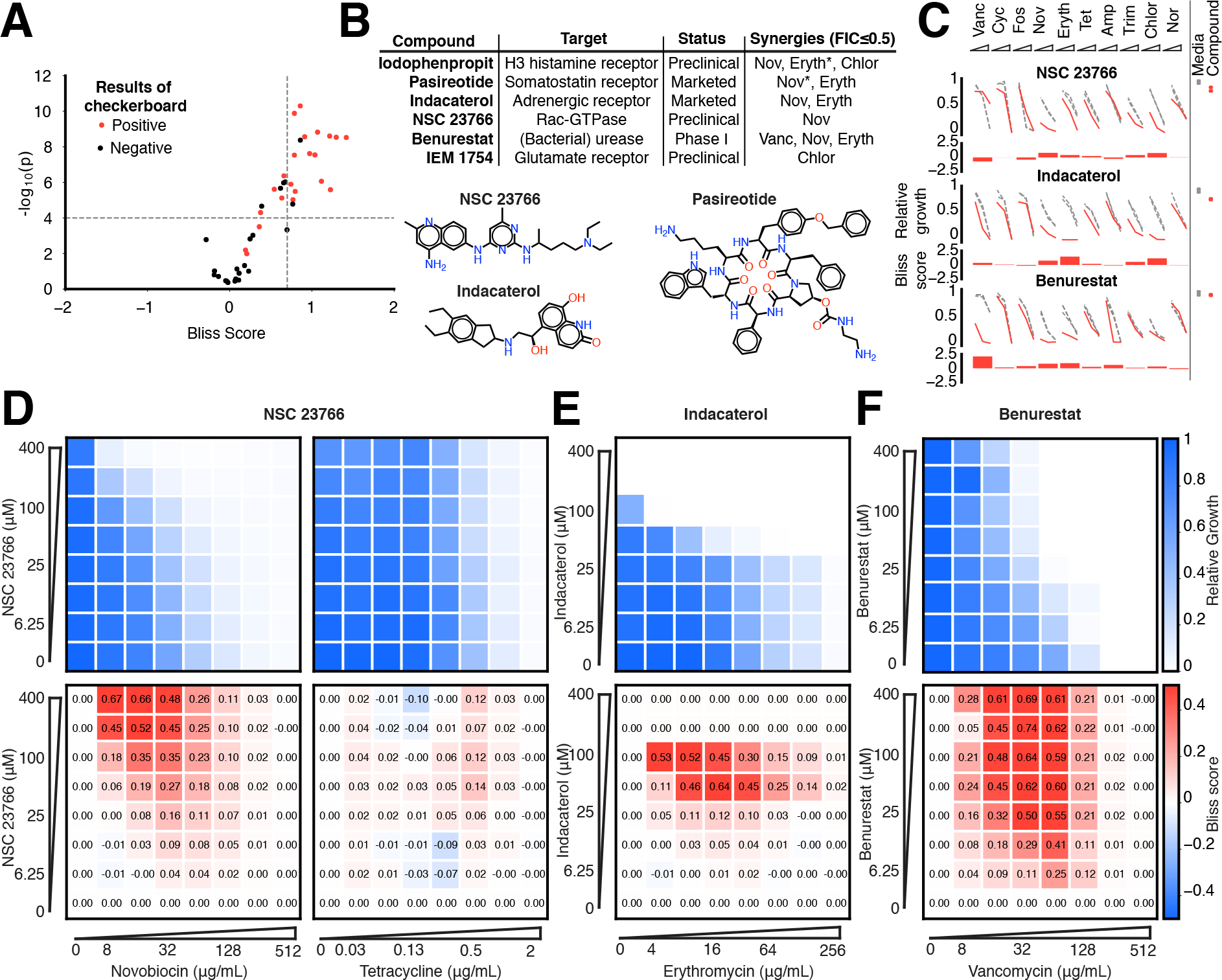
Validation of hits from primary screen. (**A**) In eight-point checkerboard assays, we tested a total of 46 compound × antibiotic combinations of which 17 combinations (11 distinct compounds) scored as hits in the primary screen (**Fig. S12**). Combinations that scored positive (red) and negative (black) for synergy by Bliss Independence in checkerboard assays performed in 96-well plate validation are plotted according to results from the primary screen. Gray lines indicate primary screening thresholds. (**B**) Target, status, antibiotic synergy set (by FIC method), and selected structures of validated hits (**Fig. S12**, **S13**, **S14**). Starred (*) antibiotics represent additional synergies revealed in checkerboard validation of hit compounds. The first four compounds listed are from the full-scale phase; the last two are from the pilot phase. Structures were rendered in ChemDraw from compound SMILES strings (**Table S4**). (**C**) Primary screening data (top panel) and calculated Bliss Scores (bottom) for three different hits (**Table S4**, **Fig. S13**). Growth in presence of compounds alone (red dots, “Compound”) relative to the absence of antibiotic and compound (gray dots, “Media”) are shown at right. (**D)** Relative growth data (top panel) and calculated Bliss Scores (bottom panel) from checkerboard assay of NSC 23766 × novobiocin (positive for synergy) (left column) and NSC 23766 × tetracycline (negative for synergy) (right column). (**E**) Checkerboard data for screening hit indacaterol × erythromycin (positive for synergy). (**F**) Checkerboard data for screening hit benurestat × vancomycin (positive for synergy).

After applying the more stringent synergy criterion of the FIC method (FIC≤0.5) to each checkerboard, we identified six compounds among our hits with synergies with at least one antibiotic by this criterion (four from full scale phase, two from pilot phase; **Fig. S12**, **S13**, **S14**, **materials and methods**). For two hit compounds, we identified synergies with additional antibiotics beyond what was detected in the primary screen, and upon further inspection we found these additional compound × antibiotic pairs scored close to the thresholds applied in the primary screen (**Fig. 4b**, **Fig. S12**). Notably, we found no previous indication of antibacterial activity for five of these six compounds, which constitute a range of different chemical structures, characteristics, and known biochemical targets (**Fig. 4b**). Comparing the primary screening data for each hit across the full panel of ten antibiotics shows some commonalities and differences that may provide clues as to mechanism of action (**Fig. 4c**, **Table S4**) *(28)*. For example, many hits shared common interactions with novobiocin and erythromycin (**Fig. 4c-e**), but some showed divergent effects with vancomycin ranging from strong synergy to strong antagonism (**Fig. 4c, f**). Here we demonstrated a nanoliter droplet combinatorial drug screening platform and applied it at scale to discover novel potentiators of antibiotics from a drug repurposing library. Our approach resolves compound crosstalk between droplets to stabilize the compound library on the timescale needed for phenotypic assays. This platform is compatible with commercially available lab equipment already present in many academic and industrial life science research facilities. Other groups have already demonstrated successful droplet-based culture of a wide range of organisms including human cell lines *(13, 18)* and we expect that our platform can be developed to support many types of phenotypic and biochemical assays. The use of optical microscopy for assay readout facilitates extension to a variety of disease-specific models and imaging assays including gene expression reporters and high content cell imaging. While much work remains, our platform represents an important new tool to leverage drug combinations for chemical biology and therapeutics discovery.

## Materials and methods

### Microwell array chip design and fabrication

Microwell array chips were designed for two sizes in AutoCAD (Autodesk) as a standard size (6.2 × 7.2 cm, 49,200 microwells; used presently in all chip screens) and a larger size (7.4 × 10 cm, 97,194 microwells, **Fig. 1c**). Each microwell consists of two circular geometries (diameter = 148.6 μm) set at 10% overlap (**Fig. S1**). We generated photomasks consisting of microwells arrayed in a hexagonal lattice with 50 μm inter-well spacing (FineLine Imaging). From these masks, we fabricated our designs to 100-120 μm feature height using photolithography on silicon wafers (Microchem SU8-2050). We embedded these wafers into custom molding jigs to create PDMS (Dow Corning Sylgard) chips (by soft lithography), of consistent thickness (1/4”) and droplet-loading slot location and size (**Fig. S1**). Chips were then coated with 1.5 μm parylene C by vapor deposition (Paratronix) to inhibit water loss from assay droplets, inhibit compound uptake, and stiffen the chip to prevent interior collapse during droplet loading.

### Cell culture preparation

We worked with (plasmid-borne) constitutive GFP-expressing strains of *Staphylococcus aureus* (chloramphenicol-resistant), *Pseudomonas aeruginosa* PAO1 (gentamicin-resistant), and *Escherichia coli* K-12 MG1655 (kanamycin-resistant). With the exception of experiments with *S. aureus*, which were conducted in LB media, all experiments and screens were conducted in cation-adjusted Mueller Hinton broth (CAMHB) (BD Difco).

For all three organisms, all experiments began with overnight cultures from glycerol stocks. Cells were initially transferred to 4 mL of media plus 30 μg/mL of the respective antibiotic to select for GFP-expressing cells. Cells were kept at 37C, 220 RPM. Shortly before an experiment onset, cells were diluted 1:1000 into antibiotic-containing media and their growth monitored. Upon reaching early log phase (OD ~ 0.05, or ~10^7^ cells/mL), cells were diluted into fresh media (no antibiotic) and normalized to a starting OD of 0.03-0.04. Droplet emulsifications into 1 nL volumes resulted in an initial count of about 10 cells per droplet. At saturation, cell density was estimated at 10^3^– 10^4^ cells per droplet.

### Fluorescence encoding

Each compound used in screening was pre-mixed with a unique ratio of three fluorescent dyes— Alexa Fluor 555, Alexa Fluor 594, and Alexa Fluor 647 (Thermo-Fisher Scientific). The following filter sets were used to detect the dye emissions: Alexa Fluor 555: Semrock SpGold-B; Alexa 594: Semrock 3FF03-575/25-25 + FF01-615/24-25; and Alexa 647: Semrock LF635-B. All ratios of the three dyes summed to a total dye concentration of 1 μM (assay final concentration). For a typical screen on the standard size array (**Microwell array chip design and fabrication**), we accepted 64 unique inputs (which produced 2,016 unique combinations) to generate ~10 replicates per combination per chip (after which we showed diminishing improvements in technical noise) (**Fig. 2i**, **Fig. 3a**, additional explanation in **Screening logistics**). Therefore, we required 64 distinct fluorescence ratios that could be identified with acceptable levels of misclassification (**Fig. S2a-b**, tested for 60 fluorescence ratios). In the current screening application, we found we did not have to remove potentially misclassified droplets in our assay, as the scoring of median GFP levels among replicate microwells was stable to outlying values. However, other more stringent applications of the droplet platform might benefit from a filtering step to remove such outliers depending on the application-specific tradeoff between the number of analyzed microwells and classification performance (**Fig. S2c**).

### Microwell array chip operation

All compounds and antibiotics were pre-mixed with fluorescence barcodes ahead of the screen. As the high concentration of compounds (200 μM prior to droplet merging) can affect the encoding dye fluorescence, we imaged the mixtures before emulsification to predict the fluorescence of the droplets and aid their later classification. In cases where shifts of the fluorescence ratios resulted in overlaps between ratios that should have been distinct, assay results encoded by these specific ratios were removed from the analysis.

Once the compounds and fluorescence barcodes were mixed, the total setup time per chip was about 30 minutes. This allowed for the overlapped setup of 18 chips/day by staggering the protocol (*i.e.* while one chip was being imaged, the next was being loaded with droplets). Each chip was first placed inside an acrylic assembly (10 min). The chip was suspended over a hydrophobic glass slide (Aquapel treated, custom cut glass from Brain Research Laboratories; 1.2 mm thickness) by plastic spacers (height = 250 μm). The chip was held in place within the assembly via its spontaneous adhesion to the top acrylic piece; the two halves of the assembly were held in place via neodymium magnets (**Fig. S1**, **Movie S1**, **S2**). Using 0% w/w surfactant oil, the gap between the chip and the glass created by the spacers was filled with oil.

Following normalization of cells in fresh media (**Cell culture preparation**), cells were added to the compound/dye mixtures. With compounds, dyes, and cells mixed appropriately, 20 μL of each mixture were emulsified into 20,000 1-nL droplets (continuous phase: fluorocarbon oil 3M Novec 7500 with 0.5-2% w/w RAN Biotech 008 FluoroSurfactant) using Bio-Rad QX200 cartridges and instrument or a custom aluminum pressure manifold (**Fig. 1a**).

Just before loading a chip, the relevant droplets were pooled (5 min) (total aqueous volume 200 μL, or ~200,000 droplets, per chip). The pooled droplets were mixed and injected into the chip, with draining oil recycled to sweep excess droplets away (5 mins) (**Fig. 1b**, **c**, **Movie S1**, **S2**). After loading was complete, the chip was washed with oil (0% w/w surfactant) to deplete residual surfactant. Spacers were gently removed to allow microwells to seal against the glass substrate; the two acrylic pieces of the assembly were then fixed with machine screws (**Fig. 1b**).

The chip was imaged at 2X magnification to identify the droplet fluorescence barcodes (12 min for standard size chip, see **Microwell array chip design and fabrication**). We merged the droplets to mix the compounds in each microwell by applying an AC electric field (4.5 MHz, 10,000-45,000-volt source underneath glass slide supplied by corona treater (Electro-Technic Products), ~10 seconds of exposure during which the tip of the corona treater was moved below the glass surface) (**Movie S3**). Due to the time associated with making/pooling droplets, loading the chip, and conducting this initial 2X magnification imaging, cells have been exposed to a given compound for 1-1.5 hours prior to droplet merging.

To allow cells to respond to the compounds present in each microwell, we incubated cells at 37C for 7 hours (without shaking), and then assayed growth by measuring constitutive GFP fluorescence with epifluorescence microscope at 2X magnification (**Fig. 1d**). GFP fluorescence reports how much the cells have grown during incubation with a dynamic range bounded by the initial count of ~10 cells/droplet and the 10^3^–10^4^ cells/droplet for saturated cultures whose growth was not inhibited.

### Antibiotic potentiation screening logistics

We performed a small pilot screen (30 chips) and a larger full-scale screen (126 chips with 108 passing a chip quality filter, **Fig. S9**). All data and performance analysis presented comes from this full-scale phase, although **Fig. 4** includes hits from both phases. Additional supporting data for pilot phase hits is shown in **Fig. S13** and **Fig. S14**.

For all screening, each chip received droplets containing a total of 64 input conditions, with 32 held constant across all chips (Set 1) and 32 that varied on each chip (Set 2) (**Fig. 3a**). Set 1 included [30 antibiotic conditions (10 antibiotics × 3 concentrations) + no cells] and [2 media-only controls + no cells]. Set 2 included [24-28 compounds (100 μM) + cells], [1 or 2 positive controls (sulbactam (20 μM) and/or erythromycin (5 μM)) + cells], and [1 or 2 negative “blank” media-only no-compound controls + cells]. Set 2 also included an additional [2 or 4 media-only controls + cells] that were used to measure dose response of antibiotics alone for comparison (**Fig. 3a**, **d**, gray dotted lines). All conditions were in CAMHB and 2% DMSO (all concentrations reported are final concentrations). We used *E. coli* K-12 MG1655 cultures (“+ cells”) with normalized starting density (**Cell culture preparation**).

We screened compounds from the Broad Institute’s Drug Repurposing Library, which consists of 4,160 compounds in 52 96-well plates (80 compounds per plate at 100 μM final concentration, with controls in columns 1 and 12). Each 96-well plate was divided into 3 groups of 32, each of which was screened on a separate chip but pooled with the same set of antibiotics-carrying droplets. This setup also gave rise to the variable number of control conditions present on each chip, as noted above.

The expected number of replicate microwells for a single chip was determined by the number of possible unique input droplet pair combinations and the number of microwells on a given chip (**Fig. S1**). For example, the standard size chip (**Microwell array design and fabrication**) had 49,200 microwells. On average, 26,772 microwells passed all quality filters. Microwells containing an [antibiotic + no cells] droplet paired with a [compound + cells] droplet constituted, on average, 10,620 of these microwells (slightly less than half due to the impact of including control combinations) (**Fig. 3c**). In order to attain an average representation of each compound × antibiotic pair of ~10 replicates (**Fig. 2i**, **Fig. 3a**), we loaded an input library of 64 unique conditions (30 antibiotic conditions + 24-28 compounds + 4-8 controls). This constituted 2,016 unique combinations, half of which (1,008) were unique compound × antibiotic combinations or controls. Overall, we achieved a median count of 13 microwells per unique compound × antibiotic combination (**Fig. 3c**).

To measure growth inhibition, we evaluated the median GFP fluorescence intensity of the replicate microwells containing a given combination as a stable statistic for the central tendency of the intensity distribution across microwells. For example, the combination [Antibiotic A + no cells] paired with [Compound B + cells] was represented in ~13 microwells on a chip (**Fig. 3c**). The median GFP intensity across these 13 replicates was used to predict whether Compound B potentiated the activity of Antibiotic A at these concentrations (**Bliss synergy scoring**). Given the nature of the chip setup, the combination [Compound B + cells] paired with [Compound C + cells] was also be represented ~13 times on this chip, enabling us to also score compound × compound synergies in addition to compound × antibiotic synergies (**Fig. S11**), though this was not the focus of our analysis.

We note that had we loaded a smaller library of, for example, 32 unique compounds (496 unique combinations), we would have attained a median representation of ~13 × 4 = 52 replicates per combination. For the noise levels associated with the present screen, this increase in replicate number would only marginally improve data quality (**Fig. 2i**) at a >4X reduction in throughput.

In summary, in the standard chip size, one chip accepted a library of 64 compounds, generating a median of 13 replicates for each of 2,016 possible combinations, half of which were either compound × antibiotic combinations or control combinations. The pilot screen (30 chips) sampled 24,000 compound × antibiotic combination assay points (800 × 30). The full-scale phase (126 chips with 108 passing a chip quality filter, **Fig. S9**) sampled 100,800 compound × antibiotic combination assay points (3,360 × 30).

### Fluorescence microscopy

All fluorescence microscopy was performed using a Nikon Ti-E inverted fluorescence microscope with fluorescence excitation by a Lumencor Sola light emitting diode illuminator (100% power setting). Images were taken across four fluorescence channels for GFP (Semrock GFP-1828A) and the three encoding dyes, Alexa Fluor 555, 594, 647 (**Fluorescence encoding**). Images were collected by a Hamamatsu ORCA-Flash 4.0 CMOS camera (exposure times range 50ms – 500ms) at 2X optical magnification (with 2X pixel binning) or 4X optical magnification (with 4X pixel binning) for 6.5 μm/pixel resolution in both cases.

### Microwell array chip image analysis

To determine the effect of each pair of input conditions, we performed a computational image analysis that (a) identified droplet pairs in each microwell; (b) assigned each droplet to an input condition using the three fluorescence colors comprising the fluorescence barcode; (c) matched the response assay signal at a later timepoint to each microwell and corresponding droplet pair; and (d) computed a statistic on all assay signals from all microwells containing the same pair of input conditions. All analysis was performed with custom Matlab and python scripts.

To detect each droplet in the image, we used a circular Hough Transform (scikit-image) *(32)* to detect circular fluorescent objects with a diameter of 100–140 μm. We inferred that a pair of droplets shared a microwell if the distance between their centroids along the vertical well axis was less than an adjustable distance threshold, typically set to 162.5 μm.

Each droplet was assigned to an input condition by determining the relative fluorescence of each of the three dyes. The three-color dye fluorescence of each droplet was projected onto a two-dimensional plane, eliminating systematic effects from differences in illumination (**Fig. S2a**). The DBSCAN algorithm (scikit-learn) *(33)* identified the clusters of droplets corresponding to each input condition, with an option for user input to correct errors, such as cluster collisions caused by optical activity of compounds in the screening library. A quality score for each droplet was computed based on the distance to the centroid of the assigned cluster (**Fig. S2b**, **c**). The Hungarian algorithm (scikit-learn) *(34)* then mapped each cluster to the pre-determined centroids of each dye mixture barcode. Pre-determined centroids were set by *a priori* ratios of dyes; or, to account for dye shifts caused by optical activity of compounds in the screening library, each input-barcode mixture was imaged prior to emulsification to predict effects on the fluorescence of the resultant droplets. Each microwell was matched with the later imaged response assay measurement (*e.g.* growth reported by constitutive GFP expression) according to the microwell position in the array (**Fig. 1d**).

### Chip quality scoring

To quality score each microwell array chip, we measured the difference between conditions representing the top and bottom of the assay dynamic range. The top of the dynamic range is given by microwells that contained a [media-only control + cells] droplet paired with a [media-only control + no cells] droplet. To represent the bottom of the dynamic range, we used microwells containing a [media-only + cells] droplet paired with a [cycloserine (16 μg/mL) + no cells] droplet. The latter condition generates a signal that represented the lowest GFP signal level expected in a growth-suppressed assay culture.

We quantified the observed dynamic range on each microwell array chip by computing the Z-factor metric *(35)*. We computed the median GFP values for the set of microwells representing the maximum and minimum signal levels described (μ_+_ and μ_–_). To estimate a standard error, we bootstrap resampled median GFP estimates from each set (1000 iterations) to estimate a sampling distribution, and measured standard errors as the standard deviation of the two sampling distributions (σ_+_ and σ_–_) (**Fig. S9**). We then computed the Z-factor (Z’) as follows:

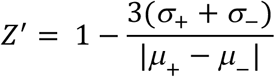

Chips with Z-factors < 0.21 were removed from analysis for low quality, a threshold determined by manual examination of the distribution (**Fig. S9b**).

### Antibiotic potentiation assay performance scoring

For each antibiotic in our panel (**Fig. 3a**, gray dotted lines), the dynamic range of the potentiation was set by the difference between relative growth measured for the lowest concentration of the antibiotic tested and maximal possible growth inhibition. To represent maximum growth inhibition in each case, we chose cycloserine (16 μg/mL), which produced the minimum detectable signal.

To quantify assay performance for the overall screen, we compiled relative growth estimates (GFP) from all chips for each antibiotic (at the lowest concentration represented) and compared to cycloserine (16 μg/mL) (**Fig. S15**). We quantified this comparison by computing the Z-factor (Z’) *(35)* for each pair of distributions across the 108 chips from the full-scale screen phase.

### Bliss synergy scoring

To estimate synergy between compounds and antibiotics, we used the deviation from growth inhibition expected by Bliss Independence *(23)*. If treatment Antibiotic A resulted in 80% growth (1-*f_A_*), and Compound B resulted in 75% growth (1–*f_B_*), then assuming they act independently we expect their combination to have resulted in 60% growth (1–*f_A_*) × (1–*f_B_*). To estimate their synergy, we subtract the observed growth from the expected growth [(1–*f_A_*) × (1–*f_B_*)]–(1–*f_AB_*) = *f_AB_* – (*f_A_+f_B_* – *f_A_ f_B_*).

We estimated the net growth inhibition for Antibiotic A from microwells carrying an [Antibiotic A + no cells] droplet paired with a [media-only + cells] droplet. We estimated net growth inhibition of a compound from microwells carrying a [Compound B + cells] droplet and a [media-only + no cells] droplet. We normalized all growth values to microwells containing a [media-only + cells] droplet paired with a [media-only + no cells] droplet and estimated synergy using the Bliss Independence metric. Since each antibiotic was present at three concentrations, we summed this metric for the compound across each of the three conditions to yield a final metric we called “Bliss Score.” Compounds with net growth inhibition exceeding 80% at the screening concentration of 100 μM were removed from analysis (**Table S5**).

We divided Bliss Scores by their corresponding standard errors to yield a test statistic (Bliss Score/standard error). To estimate the standard error in our Bliss Score measurement, we first bootstrap resampled (100 iterations) all microwells in the array to the number of replicate microwells counted for each pair of inputs. We then computed a Bootstrapped Bliss Score for each bootstrapped sample to estimate a sampling distribution, from which we computed an estimated standard error.

Using the Bliss Score and estimated standard error, we computed a test statistic (Bliss Score/standard error) that we modeled with a T-distribution fit to our blank negative controls (density function *f_T_* fit with parameters: υ = 11.23, degrees of freedom; σ = 0.922, scale parameter, fit with scipy) (**Fig. S10**).

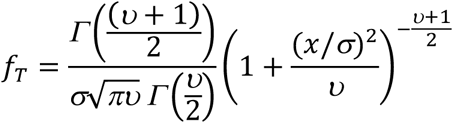

### Checkerboard validation protocol

Checkerboards were constructed from 2-fold serial dilutions of a compound and an antibiotic to create a 64-point matrix in 96-well v-bottom plates (Costar), 2% DMSO (final concentration). An *E. coli* K-12 MG1655 culture was prepared (**Cell culture preparation**) and added to the plates (final volume 100 μL; plate edges wrapped with parafilm to reduce edge effects). We incubated the plates (37C, 220 RPM, 7 hours) and measured growth by GFP accumulation using a SpectraMax plate reader (Molecular Devices) (**Fig. S12**).

We calculated deviation from Bliss Independence according to the formula above for each point in the dosage matrix (using the dosage response curves for compound alone and antibiotic alone). To compare our plate data with primary screen data, we took the maximum of Bliss Scores summed over three contiguous antibiotic doses (at the same compound dosage), for any of the top four compound doses tested. This flexibility accounted for variation in potency possibly attributable to (a) systematic differences between the 96-well plate format and our droplet platform, (b) day-to-day variation in culture conditions, and (c) the fact that all compounds were re-ordered for the validation checkerboard testing and may not have had identical formulation or purity as the compound sample used in the primary screen.

We called a compound × antibiotic pair positive for Bliss synergy in checkerboard assays if the maximum summed Bliss Score was ≥ 0.4. We computed a validation rate by computing the fraction of hit compound × antibiotic pairs in primary screening data (Bliss score > 0.7 and p-value < 10^-4^) that scored positive in the plate-based checkerboard validation testing as well, and compared this to a binomial null model with probability equal to the fraction of total positive pairs from all pairs tested.

### Fractional inhibitory concentration determination in checkerboard validation

The fractional inhibitory concentration (FIC) is a more stringent test of synergy, defining synergy as FIC ≤ 0.5 *(24)*. For a given Antibiotic A and Compound B, we measured the minimum inhibitory concentration (MIC) of each as the first well in the dosage series with <10% growth. If this was not observed in the conditions tested, we assumed that the MIC was twice the highest tested dose. For a well in the matrix at dosage point (*A: x, B: y)* with <10% growth, we calculated the FIC = *x/*MIC_A_+ *y/*MIC_B_ (where MIC_A_ is the MIC measured for Antibiotic A independently, and MIC_B_ is the MIC measured for Compound B independently). We classified the compound × antibiotic combination as synergistic if the minimum FIC in the matrix was ≤ 0.5.

### Comparison of cell growth on droplet platform with standard methods

To compare growth rates between conventional broth culture and the droplet platform, we prepared cultures of *P. aeruginosa*, *S. aureus*, and *E. coli* (**Cell culture preparation**). We then split cultures between (1) Erlenmeyer flasks (10% of the flask volume, 37C, 220 RPM) and (2) droplets loaded into the microwell array (37C, no shaking). We monitored growth via accumulation of GFP fluorescence measured by (1) transferring to clear-bottom 96-well plates and measuring by fluorescence plate reader (Molecular Devices SpectraMax), or (2) epifluorescence microscopy (Nikon Ti-E) (**Fluorescence microscopy**). For each organism, we transformed GFP measurements from the (1) 96-well plate reader to the same scale as the (2) droplet platform measurements by computing a least squares linear regression between measurements matched at each timepoint (**Fig. 2e**, data shown for (1) are transformed based on linear regression).

To compare antibiotic response curves, we created serial dilutions of six (for *P. aeruginosa*) or 12 (for *S. aureus*, *E. coli*) antibiotics in media in clear-bottom 96-well plates: Trimethoprim (Trim), Chloramphenicol (Chlor), Ceftriaxone (Ceft), Tetracycline (Tet), Kanamycin (Kan), Norfloxacin (Nor), Fosfomycin (Fos), Cycloserine (Cyc), Vancomycin (Vanc), Erythromycin (Eryth), Ampicillin (Amp), Novobiocin (Nov) (Sigma-Aldrich) (**Fig. 2f**). We emulsified cells cultured under the same conditions as above, and in parallel emulsified five points on each antibiotic dosage curve (no cells added). After pooling all emulsions, we loaded them in two technical replicate microwell arrays. Similarly, we then added cells to the 96-well plates and split the cell-antibiotic mixtures across two technical replicate plates (clear-bottom 96-well plates, with parafilm at edges to prevent edge effects, final volume 200 μL).

Using GFP fluorescence measurements similar to above, we compared the median GFP value from microwells that received an [antibiotic + no cells] droplet paired with a [media-only + cells] droplet, with equivalent final dosage conditions in the 96-well plates (**Fig. S4**, **S5**, **S6**). To compare dose responses, we obtained a non-linear least squares fit of the Hill curve for concentration *C* to data obtained from both 96-well plates and our platform (three local parameters: offset, magnitude, IC_50_; 1 global parameter for each antibiotic: Hill coefficient, *h*).

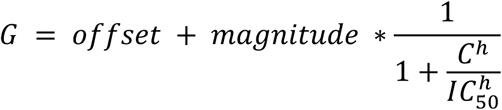

In comparing fit IC_50_’s, we removed antibiotics with plasmid-mediated resistance (*S. aureus:* Chlor; *E. coli*: Kan), or poor fit quality due to suboptimal dosage range (*S. aureus:* Ceft, Eryth, Nor, Tet; *E. coli:* Ceft).

As construction of droplet pairs in microwells was stochastic, each pair of input conditions appeared in a random number of replicate microwells. To compute how technical noise scales as a function of *k* replicate microwells, we resampled with replacement *k* microwells from each set of replicates and recomputed the median GFP across each sample (**Fig. 2i**).

### Assessment of cross-contamination between microwells

To construct source droplets, we emulsified resorufin (10 μM) in CAMHB using 2% w/w 008 FluoroSurfactant (RAN Biotech) in Novec 7500 fluorocarbon oil (3M). Sink droplets were made in a similar fashion, but with fluorescein (5 μM, CAMHB), as this dye showed negligible exchange on assay timescales *(20, 21)*. Droplets were pooled in 1:1 ratio (5 min) and then loaded into the chip (5 min) such that microwells received either two sink droplets (sink-only wells), two source droplets (source-only wells), or one sink and one source droplet (**Fig. 2a**). The random loading process resulted in these pair types being randomly dispersed across the array. We washed the chip with oil containing either 2%, 0.5%, or 0% w/w surfactant, and then mechanically clamped the chip to the glass substrate (**Fig. S1**) according to our standard screening protocol.

Resorufin fluorescence measurements of the sealed array were taken over 20 hours at three distinct fields of view. Sink droplets were identified by their high fluorescein fluorescence, and assay background was subtracted from all measurements. To measure inter-well exchange, we compared the mean resorufin fluorescence of microwells containing two source droplets and microwells containing two sink droplets (normalized to their sum) (**Fig. 2b-d**). As a proxy for bulk emulsion systems and to predict exchange during the pooling phase of our protocol, we compared the mean resorufin fluorescence of source droplets and sink droplets in microwells containing one source droplet and one sink droplet (normalized to their sum) (**Fig. 2b-d**, **Fig. S3a-b**).

To measure the relationship of exchange kinetics and number of neighboring droplets, we additionally created arrays with microwells sized to receive seven droplets. This microwell array chip was washed with 2% w/w surfactant, and all other experimental conditions were the same as above (**Fig. S3c-e**).

## Supplementary Text

### Estimation of compound exchange kinetics during pooling phase

Compartmentalization of droplets in the microwell array chip and depletion of free surfactant limits compound exchange between wells (**Fig. 2a-d**). However, limited quantities of compounds may still exchange between droplets via supra-molecular surfactant complexes such as reverse-micelles in the continuous phase during the time that droplets are pooled prior to washing and sealing the array (**Fig. 1a**; 10 min). Reverse-micelles in the bulk fluorous oil have a fluorous exterior and a PEG interior. We hypothesize that exchange of small hydrophobic solutes occurs by partitioning of compounds from the aqueous droplet interior to the PEG phase of reverse-micelles, the dynamic formation and fusion of reverse-micelles with droplets, and the diffusion of compound-laden reverse-micelles through the continuous oil phase *(20, 21)*.

The opportunity for exchange occurs in our workflow because droplets carrying different compounds are randomly dispersed in a three-dimensionally packed bulk emulsion during the droplet pooling and mixing step, possibly allowing transport to neighboring droplets. To estimate the transport kinetics, we (i) measured the exchange rate between neighboring droplets (**Fig. S3a,b**); (ii) measured the dependence on number of neighboring droplets (**Fig. S3c-e**); and (iii) compared predictions to experimentally observed quantities (**Fig. S3f**).

As a baseline, we measured the kinetics of resorufin exchange between neighboring droplets in microwells containing one source and one sink droplet (**Fig. 2a, Fig. S3a**). We modeled exchange at early timepoints (< 2 hours) by a single-exponential (**Fig. S3b**), from which we could fit a kinetic constant (*k*) as a function of surfactant concentration *(20)*.

During droplet pooling, droplets are dispersed in a three-dimensional bulk emulsion of packed spheres, so the compound exchange rate could be increased relative to the estimate from microwells containing only two droplets as each neighboring droplet in the bulk emulsion can participate in compound exchange *(20)*. For example, a droplet carrying Compound B may neighbor two droplets carrying Compound C. To measure the relationship of neighboring droplet number and exchange kinetics, we constructed microwells that hold a total of seven droplets each, with random loading of source (resorufin) or sink (fluorescein) droplets (**Fig. S3c-d**). We found a linear relationship between exchange rate (between source and sink droplets that shared the same microwell) and the number of source droplets in the microwell (and therefore, the interfacial surface area available for surfactant-dependent exchange) (**Fig. S3c-e**). To predict compound exchange in a bulk emulsion, we linearly extrapolated the fit kinetic constants in measured in **Fig. S3b** by the number of neighboring droplets carrying a given compound (**Fig. S3f**).

As an experimental test of our prediction, we estimated the exchange between source and sink droplets during pooling in the experiment described in **Figure 2a**, **d**. Since we do not detect any increase in resorufin in microwells containing only sink droplets (**Fig. 2d**), the fraction of fluorescence detected at the first timepoint (.096) must have occurred during the pooling step. Source and sink droplets were pooled in a 1:1 ratio, so we expect that during pooling, for a given sink droplet, an average of 50% of the 8-12 neighboring droplets in the three-dimensional bulk emulsion are source droplets (with the distribution as binomial). This measurement (0.096) is in good agreement with our predictions of exchange rate for four neighboring source droplets over the duration of the pooling step (**Fig. S3f**).

Under the conditions in the antibiotic potentiation screen, we pooled droplets carrying 64 distinct inputs for each chip, so the mixing ratio was much lower than 1:1 (1:64). We expect that in these circumstances, it was rare that a given droplet neighbored more than one droplet carrying the same compound, so kinetics for exchange of any single compound are described by one neighbor (**Fig. S3f,** blue curve) and are even more limited than in the above tests with resorufin (where pooling was in a 1:1 ratio).

However, while resorufin dye is a convenient model, exchange kinetics also depend on compound properties, specifically those that affect the relative affinity of a given compound for reverse-micelles *(21)*. Empirically, we see that more hydrophobic compounds exchange faster (fluorescein exchanges more slowly, and rhodamine more quickly than resorufin) and that LogD (log_10_ of the octanol-buffer partition coefficient) is a useful predictor of exchange rate. The cLogP value (a calculated prediction of hydrophobicity) of resorufin is 1.77, which is of middling hydrophobicity compared with the compounds used in our screen.

There is ample evidence that the droplet platform performs adequately despite the potential for false-negatives (due compound loss from a droplet), and false-positives (due to exchange among droplets during the initial pooling step) by compound exchange. First, the antibiotic IC_50_’s measured in the chip corresponded closely to those measured in 96-well plates. Second, 108/124 chips without logistical failures passed stringent quality control assessments of the internal positive and control conditions. Finally, of our primary screening hits where validation was attempted, 88% (15/17 tests, p-value = 0.00058 for a binomial null model across all pairs positive in plates; 10/11 distinct compounds) were successfully validated in conventional 96-well plate checkerboard assays (**Fig. 4a**, **Fig. S12**, **materials and methods**).

---

**Figure S1.**
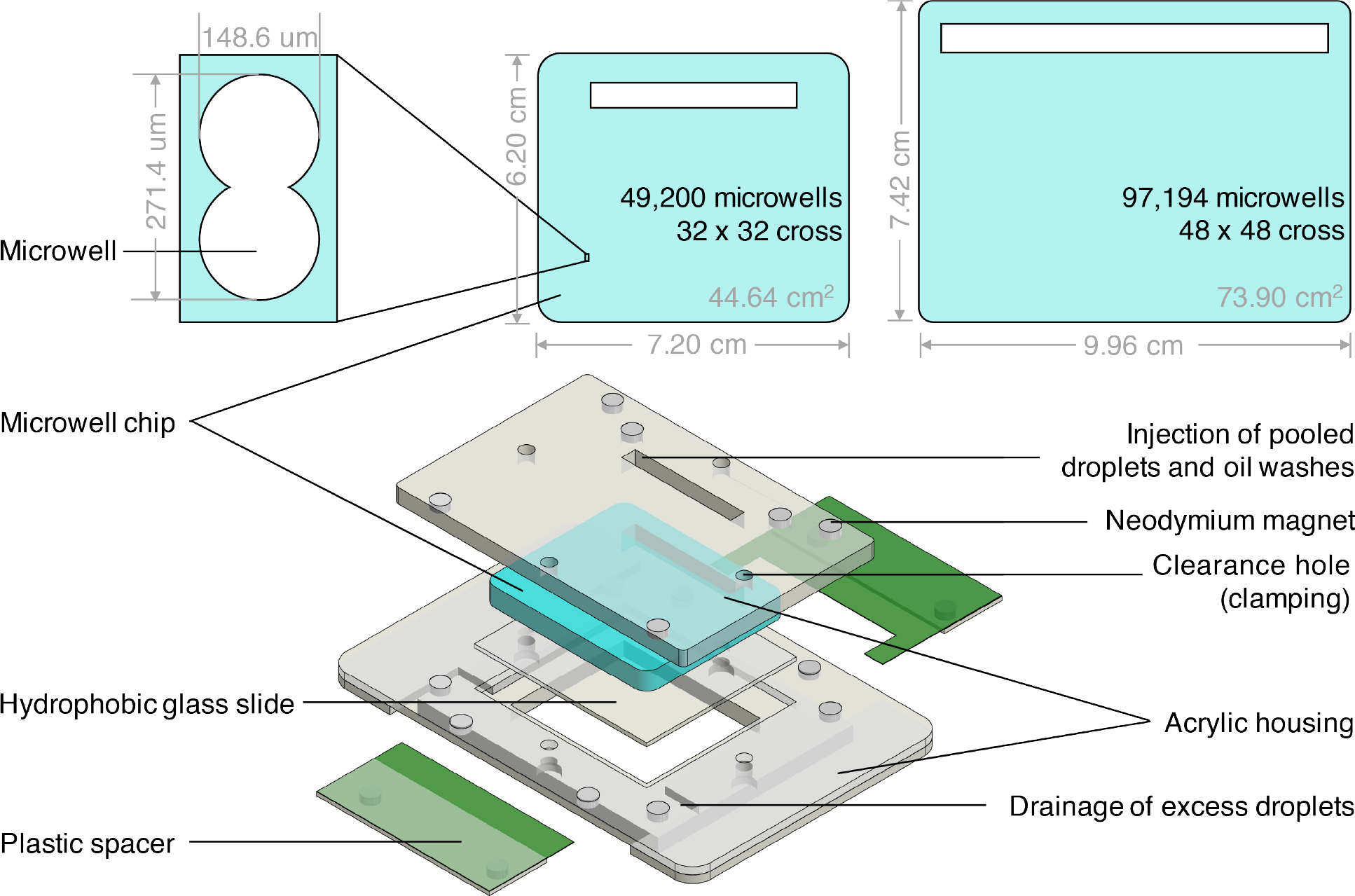
Microwell array chip and chip-loading assembly designs. We created two formats for microwell array designs of different sizes (standard size = 49,200 wells; large size = 97,194 wells). All data included in this work were collected with the standard size. The large size is represented in **Fig. 1c** and **Movie S1**, **S2**. An assembly of an acrylic clamp and plastic spacers suspends the microwell array above a hydrophobic glass slide to create a wide flow cell in which droplets are loaded. To seal the array against the glass after loading, the spacers are removed, and the acrylic assembly is secured with machine screws (**materials and methods**).

**Figure S2.**
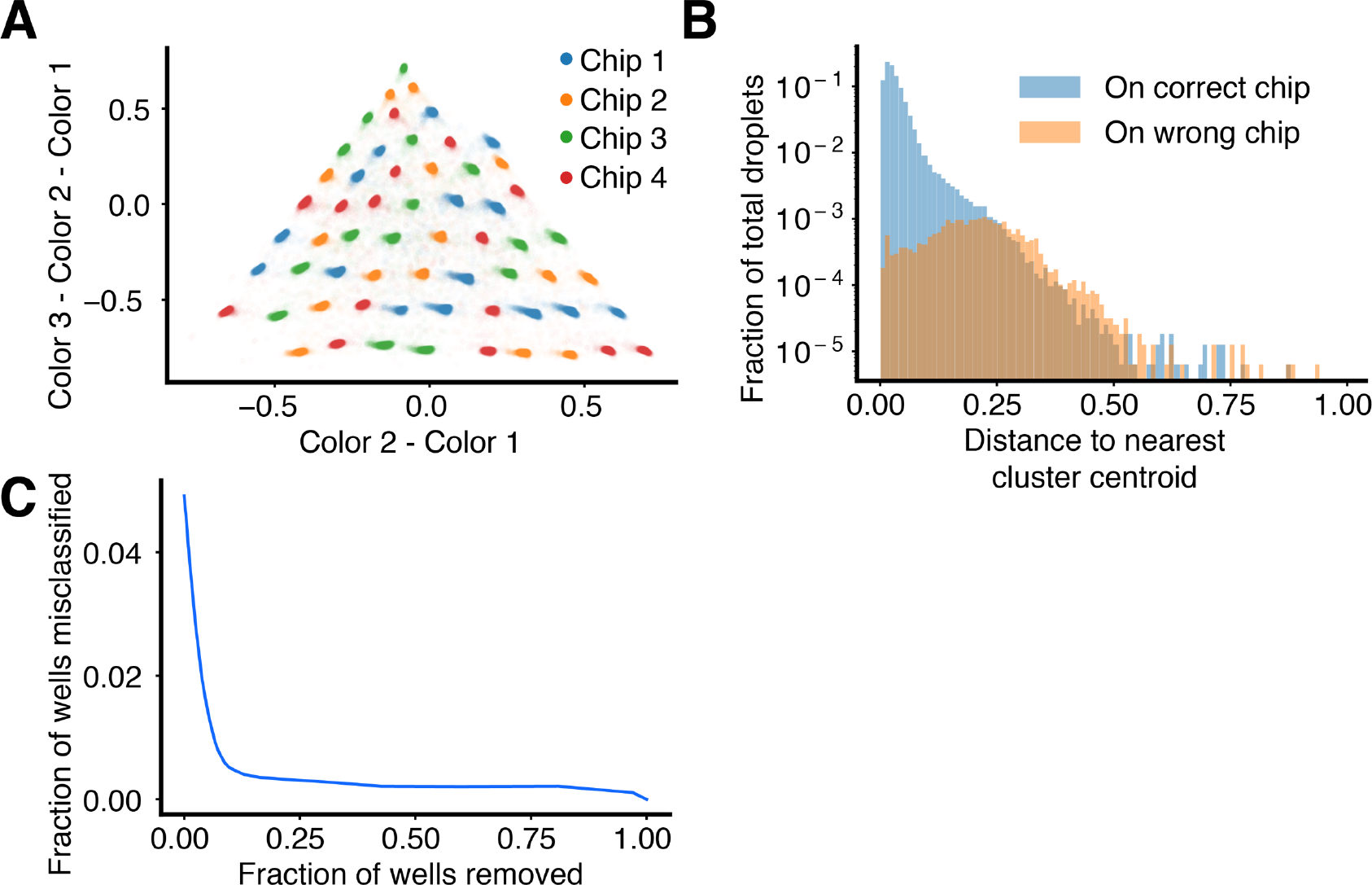
Fluorescence barcoding performance. (**A**) To measure barcode classification performance, we created droplets from a set of 60 barcodes with different ratios of 3 fluorescent dyes (Color 1 = Alexa Fluor 555, Color 2 = Alexa Fluor 647, Color 3 =Alexa Fluor 594). The set of 60 was divided into 4 sets of 15 and split across different chips (blue, orange, green, or red). The three-color fluorescence values of each droplet (n = 164,024 across 82,012 wells) are shown as a 2-dimensional projection. Each point is plotted at 0.5% opacity. (**B**) Droplets were determined to be misclassified if they were assigned to a barcode that was not represented on the chip. A histogram shows the fraction of total droplets that were misclassified as function of each droplet’s distance to its assigned barcode centroid. (**C**) Microwells with at least one droplet exceeding a distance threshold can be removed to improve classification performance. We did not find this necessary in our screening applications due to the stability of our summary statistic (median of GFP levels in replicate microwells) to outliers (**materials and methods**). Here, the fraction of wells misclassified is estimated by multiplying the number of wells with at least 1 misclassified droplet by 4/3 (60/45), since misclassification can only be detected by comparing the 15 colors present on a given chip to the 45 present on the others.

**Figure S3.**
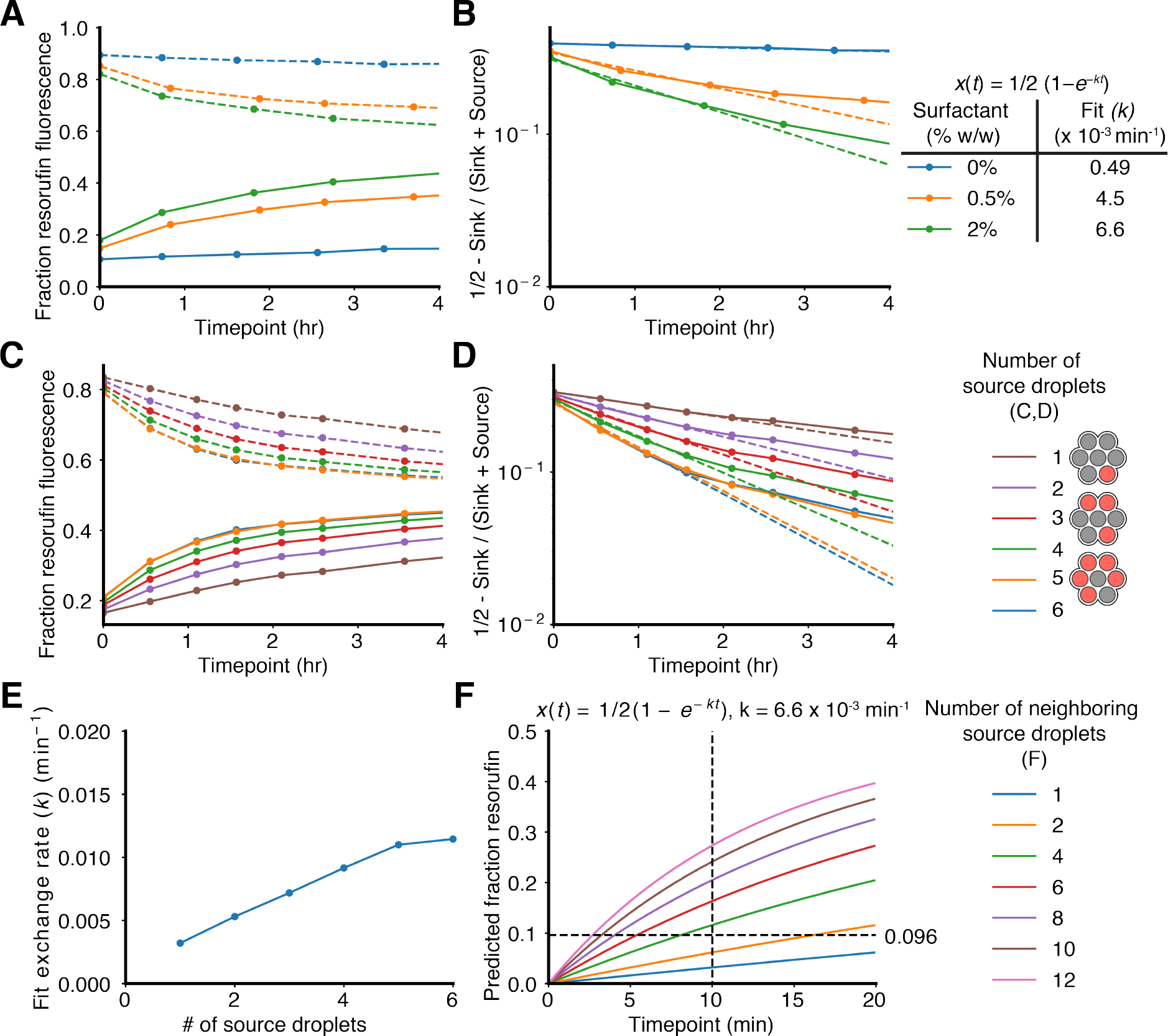
Expected exchange kinetics during pooling phase. (**A**) Exchange of the fluorescent dye resorufin between neighboring droplets can be monitored in wells containing a single source and sink droplet (**Fig. 2a**) by tracking the accumulation of dye in the sink droplets (solid) and the depletion of dye from the source droplets (dashed). Data was collected from three fields of view per chip (2% surfactant: 298 microwells analyzed; 0.5% surfactant: 441 microwells analyzed; 0% surfactant: 439 microwells analyzed). (**B**) Kinetics of exchange exhibit a single exponential at early timepoints, and a power law at later timepoints *(20)*. Exchange kinetic constants (*k*) for each concentration of surfactant were estimated from data from the first 2 hours. (**C**) During the pooling phase (**Fig. 1a**), droplets form a three-dimensional bulk emulsion that increases the transport surface area. To measure the relationship of surface area and exchange kinetics, we constructed wells that hold seven droplets total, with stochastic loading of source and sink droplets. We then measured the exchange kinetics between source and sink droplets that shared the same well, as a function of the number of source droplets. This experiment was conducted with 2% w/w surfactant. Data was collected from three fields of view (146 microwells analyzed). (**D**) We estimated exchange kinetic constants (*k*) as a function of the number of source droplets by fitting a single exponential to the first two hours. (**E**) The exchange kinetic constants show a linear relationship with the number of source droplets. (**F**) We predict the exchange during pooling by linearly extrapolating the fit kinetic constants in (**B**) by the number of source droplets neighboring a given sink droplet in a three-dimensional bulk emulsion. In the experiment described in **Fig. 2a**, source and sink droplets were pooled in a 1:1 ratio for 10 minutes. At the first timepoint, the fraction of resorufin dye in the sink droplets in sink-only wells (0% w/w surfactant) was measured as 0.096 (**Fig. 2d**).

**Figure S4.**
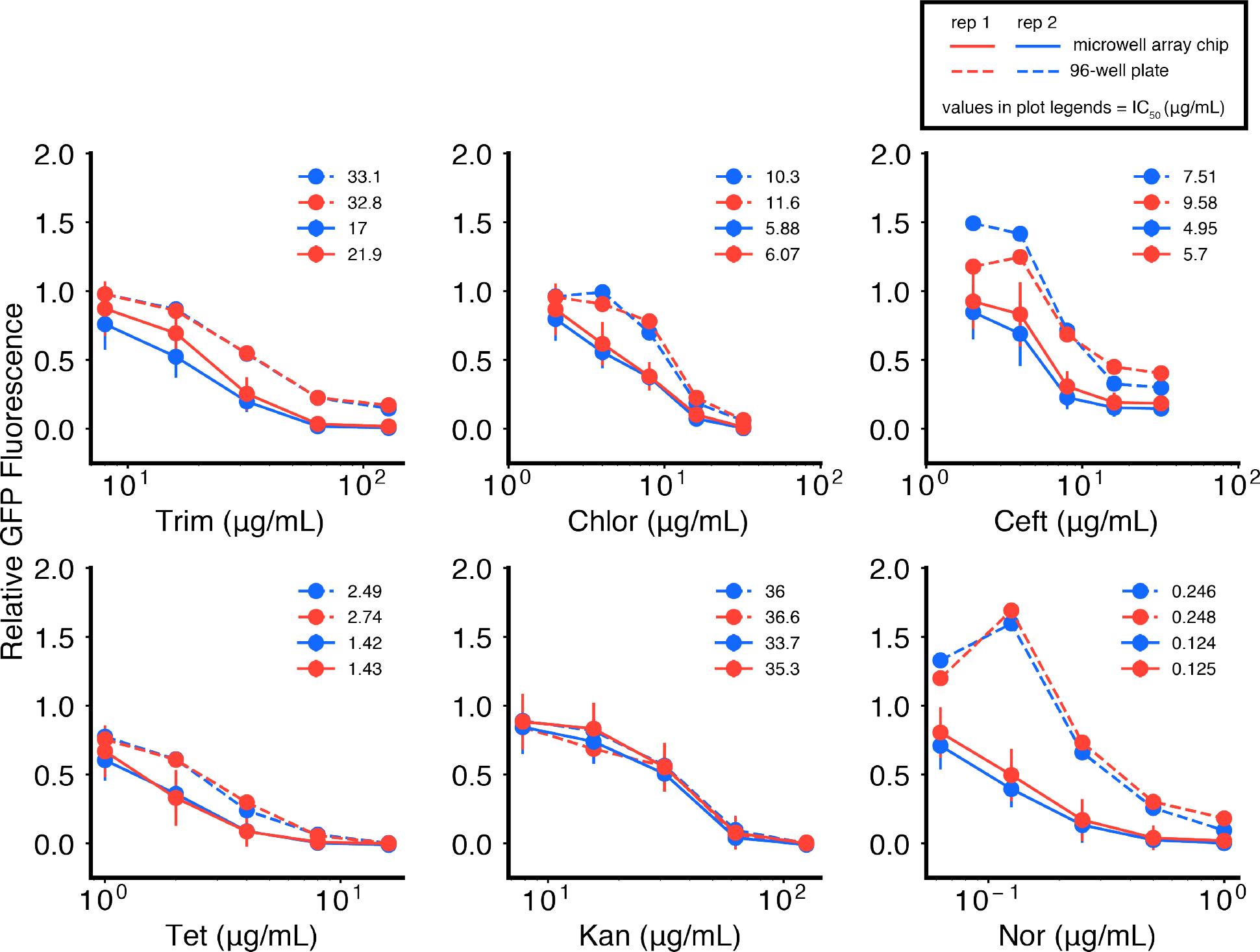
Antibiotic dose response comparison between 96-well plates and the microwell array chip for *P. aeruginosa*. We tested five-point dose response curves for 6 different antibiotics against *P. aeruginosa* and compared dose responses in 96-well plates (dotted lines) with the microwell array chip (solid lines), for two technical replicates (replicate 1: blue, replicate 2: red). All data are normalized to no-antibiotic controls. Error bars represent standard deviations of replicate microwells for each condition, with samples of size n = 127 (100, 147.5) (replicate 1), and n = 172 (149.5, 193) (replicate 2) (median (25^th^ percentile, 75^th^ percentile)). Legends represent fit IC_50_ (μg/mL) values for each curve, obtained by non-linear least squares fitting of the Hill curve to each dose response. Data points and IC_50_ values are also reported in **Fig. 2g**, **h**, **j**, and **Fig. S7**.

**Figure S5.**
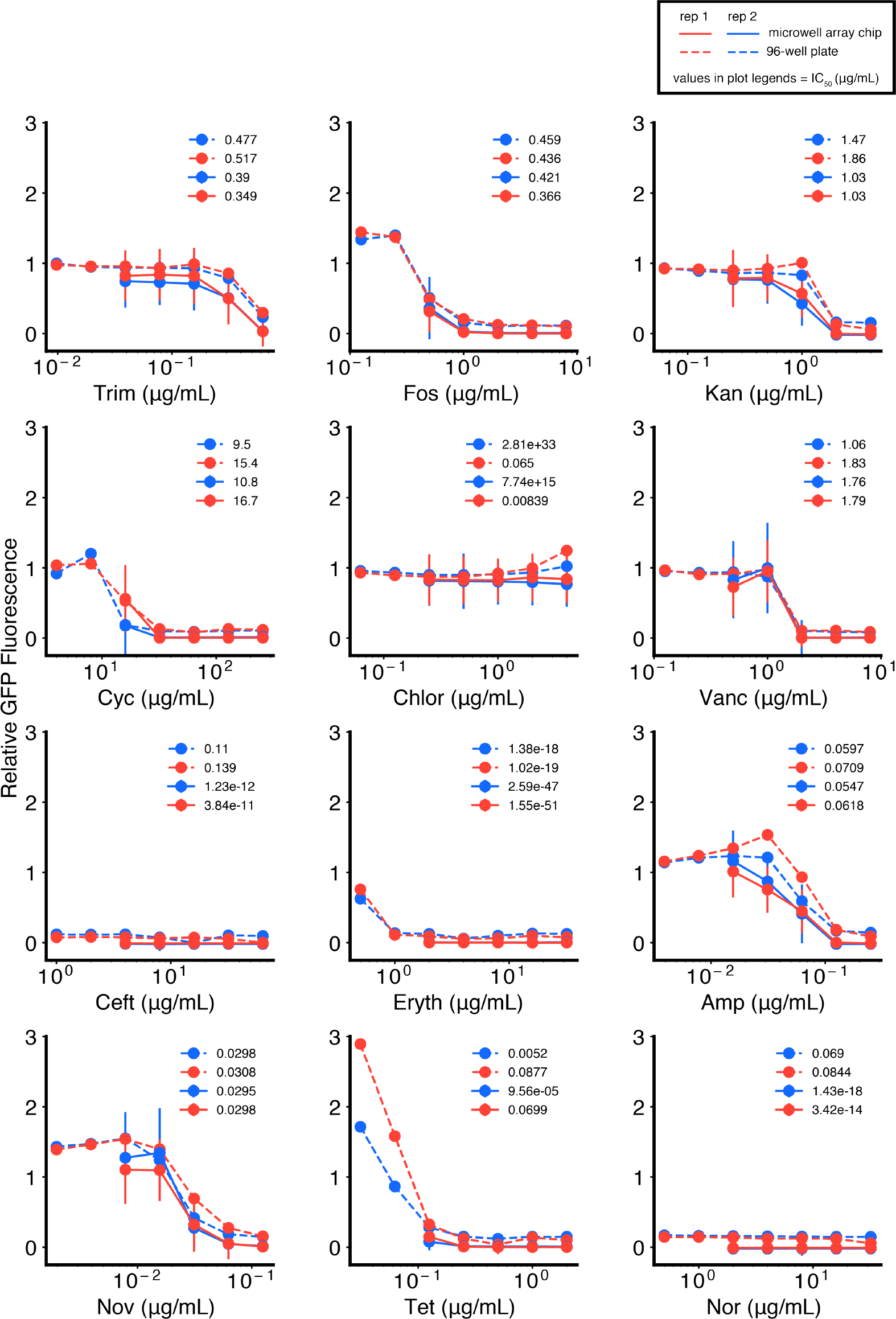
Antibiotic dose response comparison between 96-well plates and the microwell array chip for *S. aureus*. We tested five-point dose response curves for 12 different antibiotics against *S. aureus* and compared responses in 96-well plates (dotted lines) with the microwell array platform (solid lines), for two technical replicates (replicate 1: blue, replicate 2: red). All data are normalized to no-antibiotic controls. Error bars represent standard deviations of replicate microwells for each condition, with samples of size n = 157.5 (120.25, 190) (replicate 1), and n = 142.5 (119, 165.5) (replicate 2) (median (25^th^ percentile, 75^th^percentile)). Legends represent fit IC_50_ (μg/mL) values for each curve, obtained by non-linear least squares fitting of the Hill curve to each dose response. Data points and IC_50_ values are also reported in **Fig. 2g**, **h**, **j**, and **Fig. S7**.

**Figure S6.**
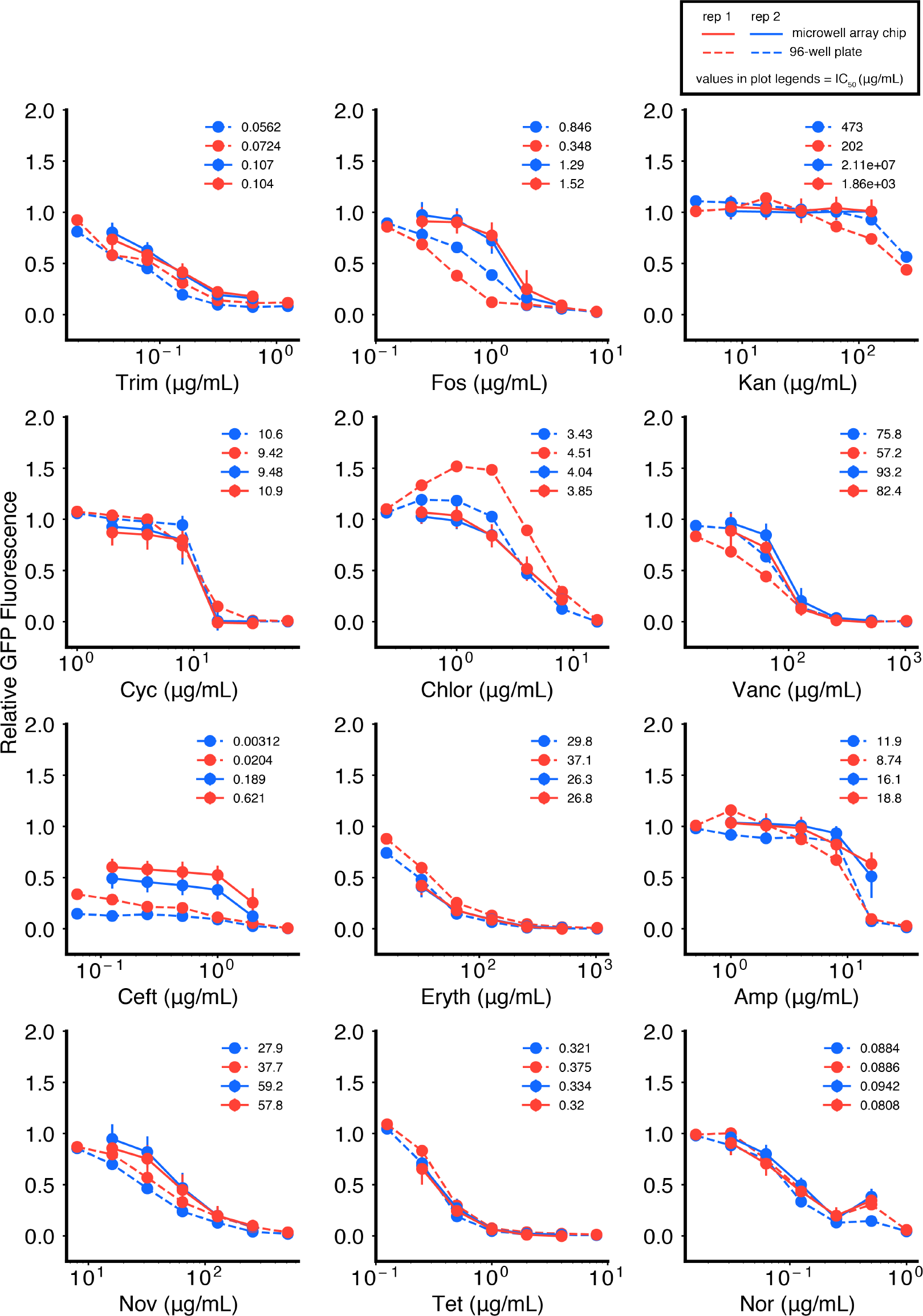
Antibiotic dose response comparison between 96-well plates and the microwell array chip for *E. coli*. We tested five-point dose response curves for 12 different antibiotics against *E. coli* and compared responses in 96-well plates (dotted lines) with the microwell array platform (solid lines), for two technical replicates (replicate 1: blue, replicate 2: red). All data are normalized to no-antibiotic controls. Error bars represent standard deviations of replicate microwells for each condition, with samples of size n = 171 (137.25, 208.25) (replicate 1), and n = 130 (107.5, 149) (replicate 2) (median (25^th^ percentile, 75^th^ percentile)). Legends represent fit IC_50_ (μg/mL) values for each curve, obtained by non-linear least squares fitting of the Hill curve to each dose response. Data points and IC_50_ values are also reported in **Fig. 2g**, **h**, **j**, and **Fig. S7**.

**Figure S7.**
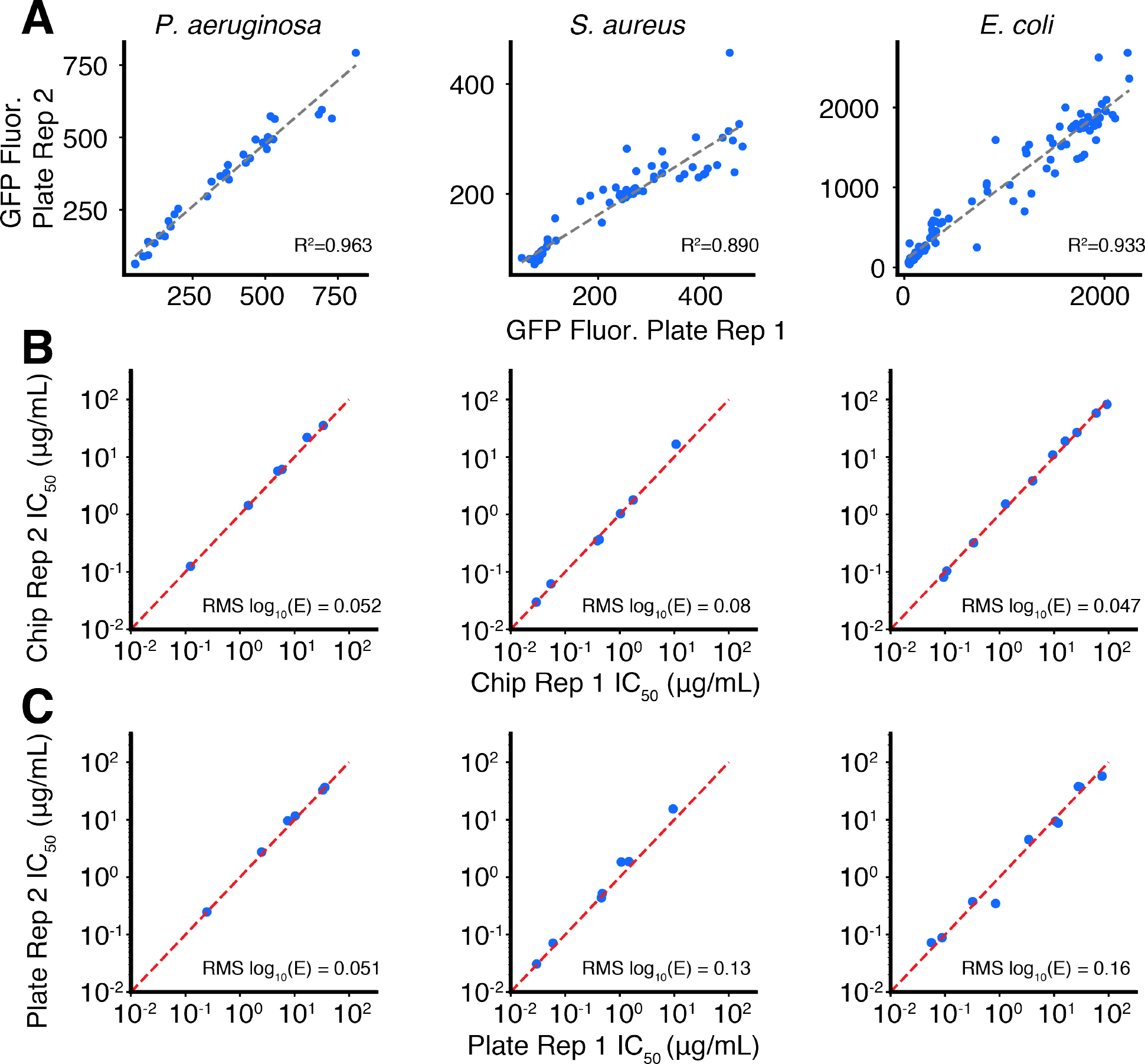
Technical noise estimation of microwell array chip and 96-well plate assays. (**A**) Scatterplot comparison of 96-well plate technical replicate measurements of relative GFP fluorescence, from **Fig. S4**, **S5**, **S6**. Lines of best fit (gray, dotted) and corresponding R^2^ values are shown. R^2^ values obtained are also shown for comparison in **Fig. 2i** (gray, dotted) and reported in **Fig. 2j**. (**B**, **C**) Scatterplot comparisons of fit IC_50_ values from technical replicate antibiotic dose responses (**Fig. S4**, **S5**, **S6**). We report the root mean square of the change in log_10_(IC_50_) [RMS Δlog_10_(IC_50_)], which represents the difference from the diagonal (*x* = *y*, red, dotted). These values are also reported in **Fig. 2j**.

**Figure S8.**
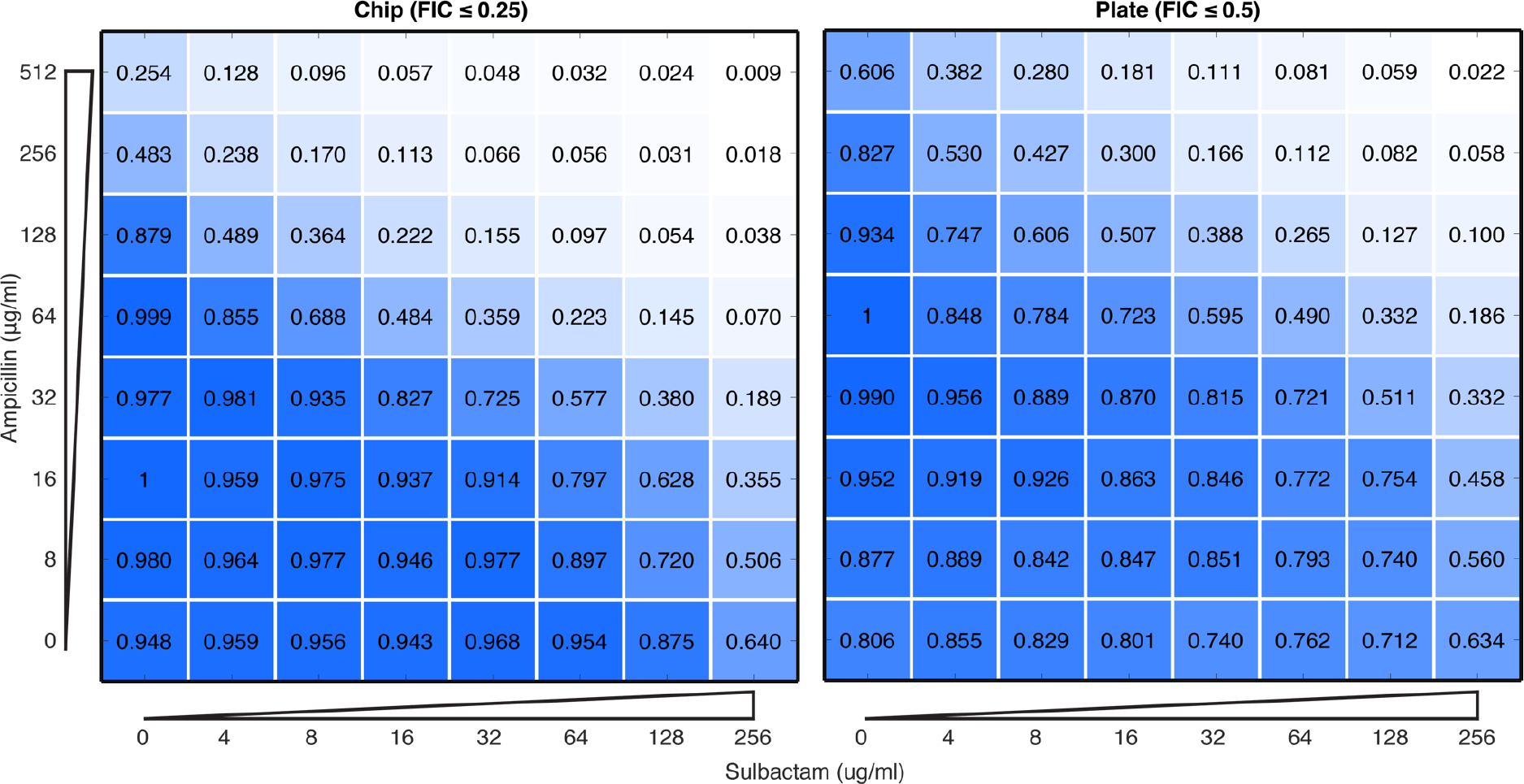
Comparison of checkerboard assay between 96-well plates and microwell array chip. Comparison of the checkerboard drug interaction assay in *P. aeruginosa* for the antibiotic ampicillin and sulbactam, a beta-lactamase inhibitor and known potentiator. Each point shows a relative growth value normalized to the maximum growth value on the 64-point matrix. Both platforms report a synergistic interaction as described by the FIC method (FIC ≤ 0.5) (**materials and methods**).

**Figure S9.**
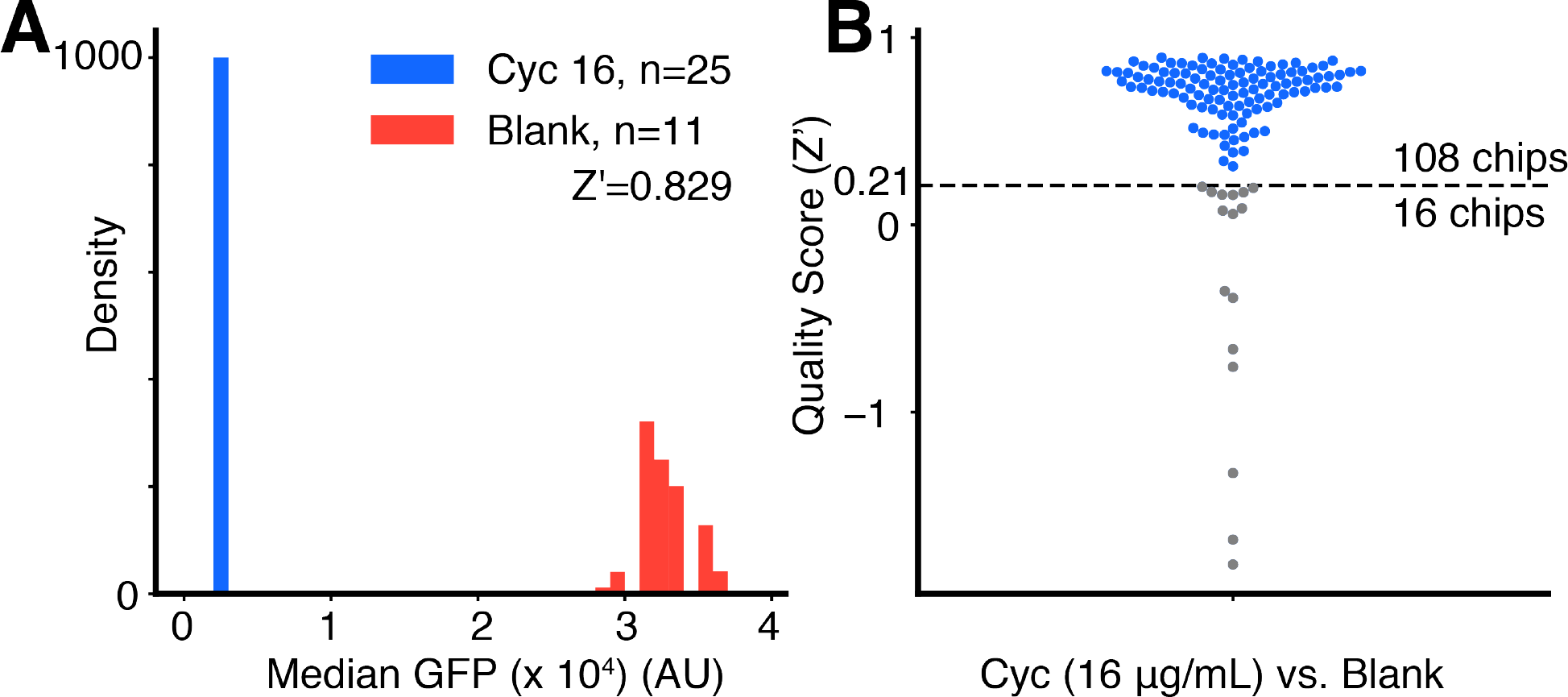
Quality scoring individual microwell array chips used in the screen. We estimate a quality score of each microwell array chip by evaluating the observed dynamic range in growth values. Chips with low quality were then filtered out prior to analysis of screening data. (**A**) Quality scoring procedure for a single representative microwell array chip based on calculating a Z-factor (Z’) between microwells representing the top (media-only control, red) and bottom (cycloserine 16 μg/mL, blue) of the growth assay dynamic range (**materials and methods**). Histograms show median GFP growth values computed from samples bootstrapped from original sample of replicate microwells (1,000 iterations). (**B**) Each point represents the Z-factor (Z’) computed for a particular microwell array chip, as shown for an exemplar chip in part A. 108 chips passed our quality threshold (Z’ > 0.21), and 16 chips failed.

**Figure S10.**
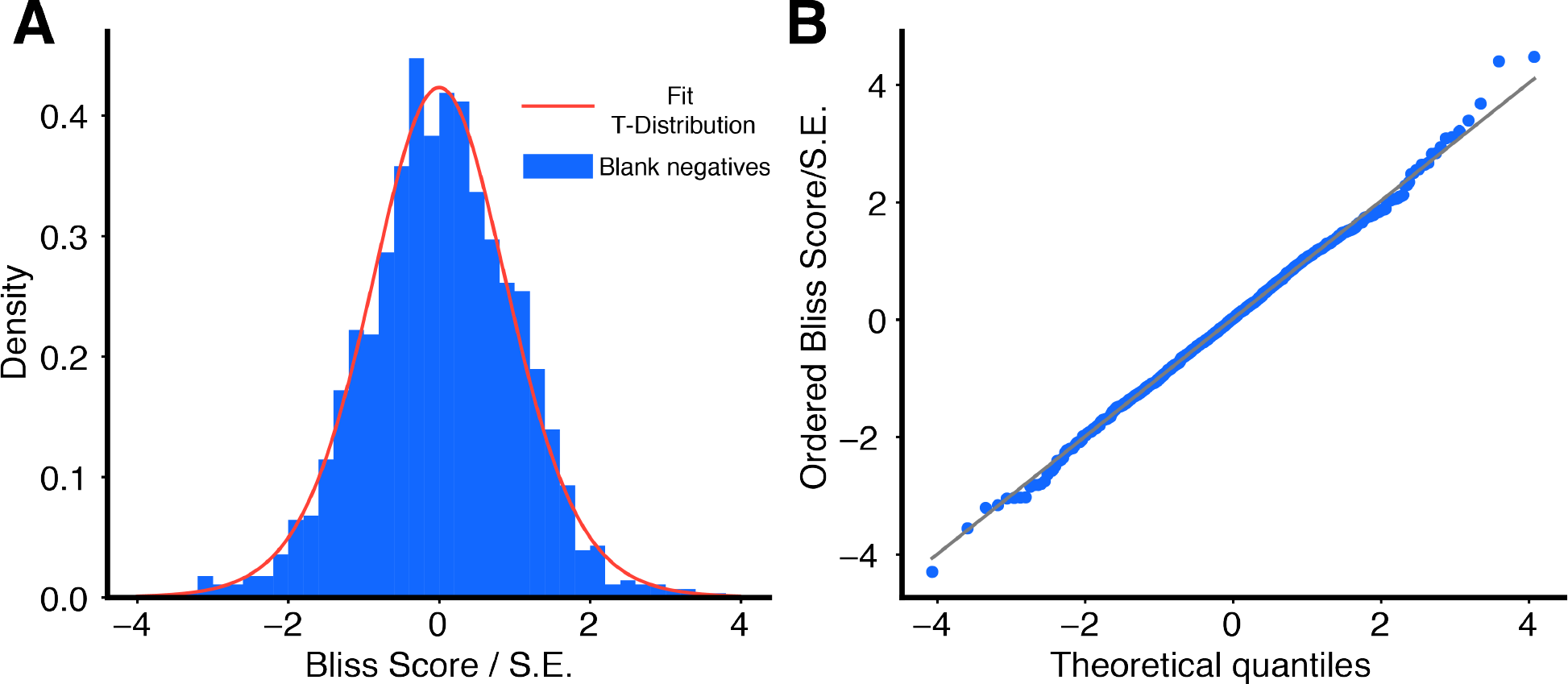
Distribution of screening blank negative controls. To estimate the statistical significance of Bliss Scores measured for each compound × antibiotic pair in the primary screen, we computed a test statistic from each Bliss Score divided by an estimated standard error (**materials and methods**). Null data representing this test-statistic were obtained in-line from the screen from all pairs of blank negative controls × antibiotics. (**A**) A histogram of the test statistic for all blank negative control × antibiotic pairs (n = 140 × 10 antibiotics = 1400; blue) and a fit T-distribution (11.2 degrees of freedom, scale = 0.992; red) which constituted our null model for p-value calculations. (**B**) To determine the quality of the fit, we compare ordered values of our test-statistic with theoretical quantiles estimated for our fit T-distribution. Theoretical quantiles are generated by Filliben’s estimate of the median order statistic *(36)*. The diagonal (*x* = *y*, gray) is shown for comparison.

**Figure S11.**
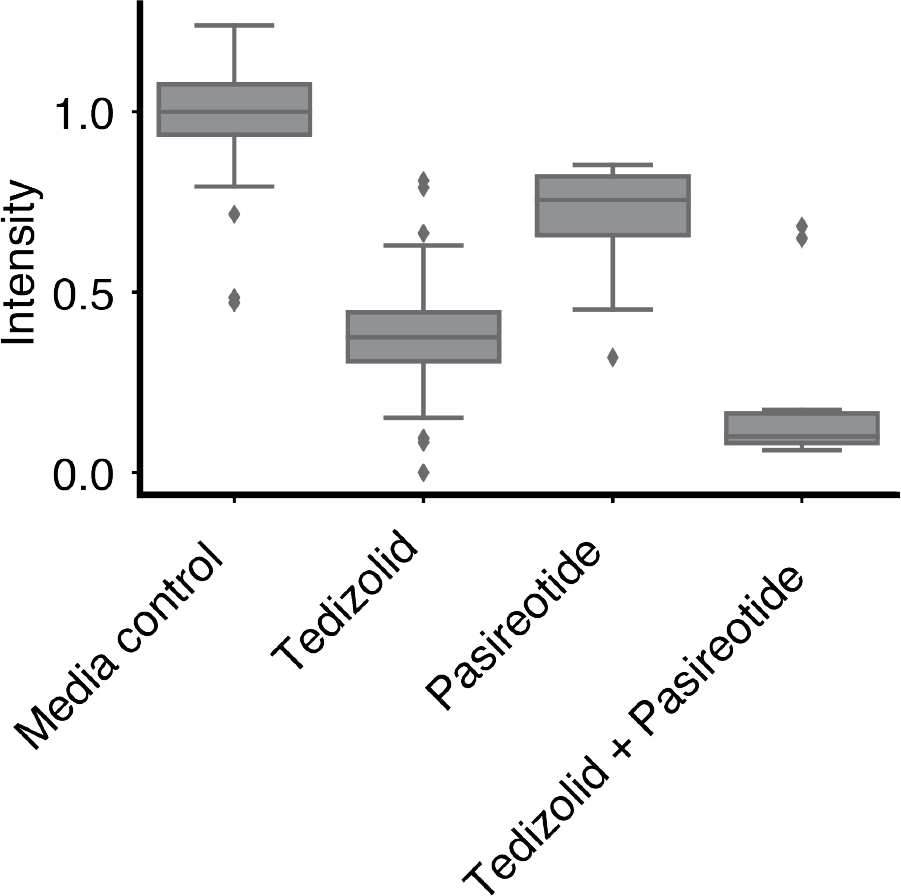
Synergy between two compounds in antibiotic potentiation screen. Tedizolid and pasireotide are two compounds from the drug repurposing library that were loaded on the same microwell array. As the array constructed compound × compound pairs in addition to compound × antibiotic pairs, we identified a synergistic interaction between these two compounds. Intensity reflects GFP measurements for each microwell, normalized to the median GFP value for microwells containing a [media-only + cells] droplet paired with and [media-only + no cells] droplet (Tukey box plot). Tedizolid reflects relative GFP measurements for microwells carrying a [tedizolid (100 μM) + cells] droplet paired with a [media-only + no cells] droplet. Pasireotide reflects relative GFP measurements for microwells carrying a [pasireotide (100 μM) + cells] droplet paired with a [media-only + no cells] droplet. Tedizolid + Pasireotide reflects relative GFP measurements for microwells carrying a [tedizolid (100 μM) + cells] droplet paired with a [pasireotide (100 μM) + cells] droplet. (Note that synergy is still detected for Tedizolid + Pasireotide despite there being an initial starting cell count that is twice as large as that of the other pairs.)

**Figure S12.**
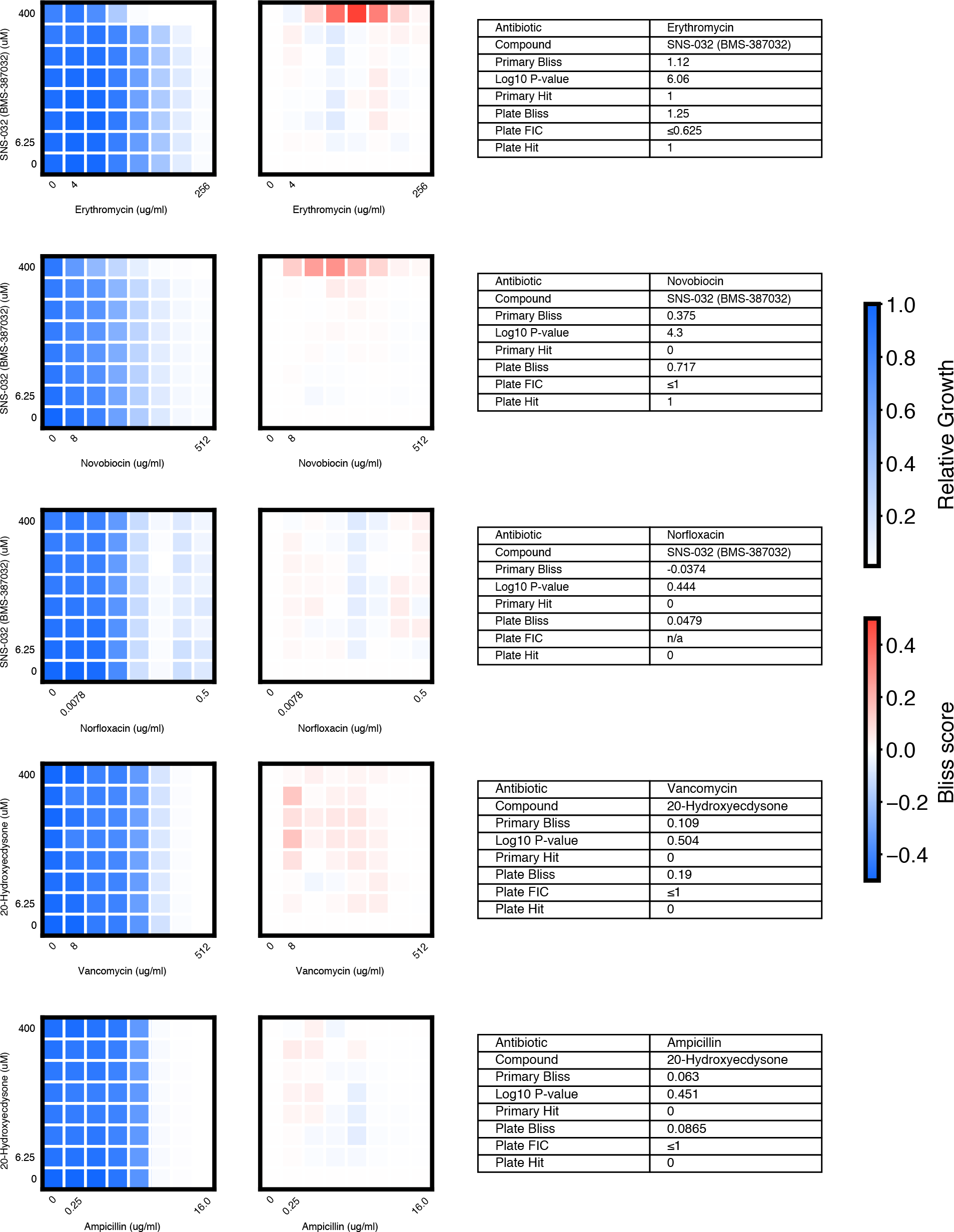
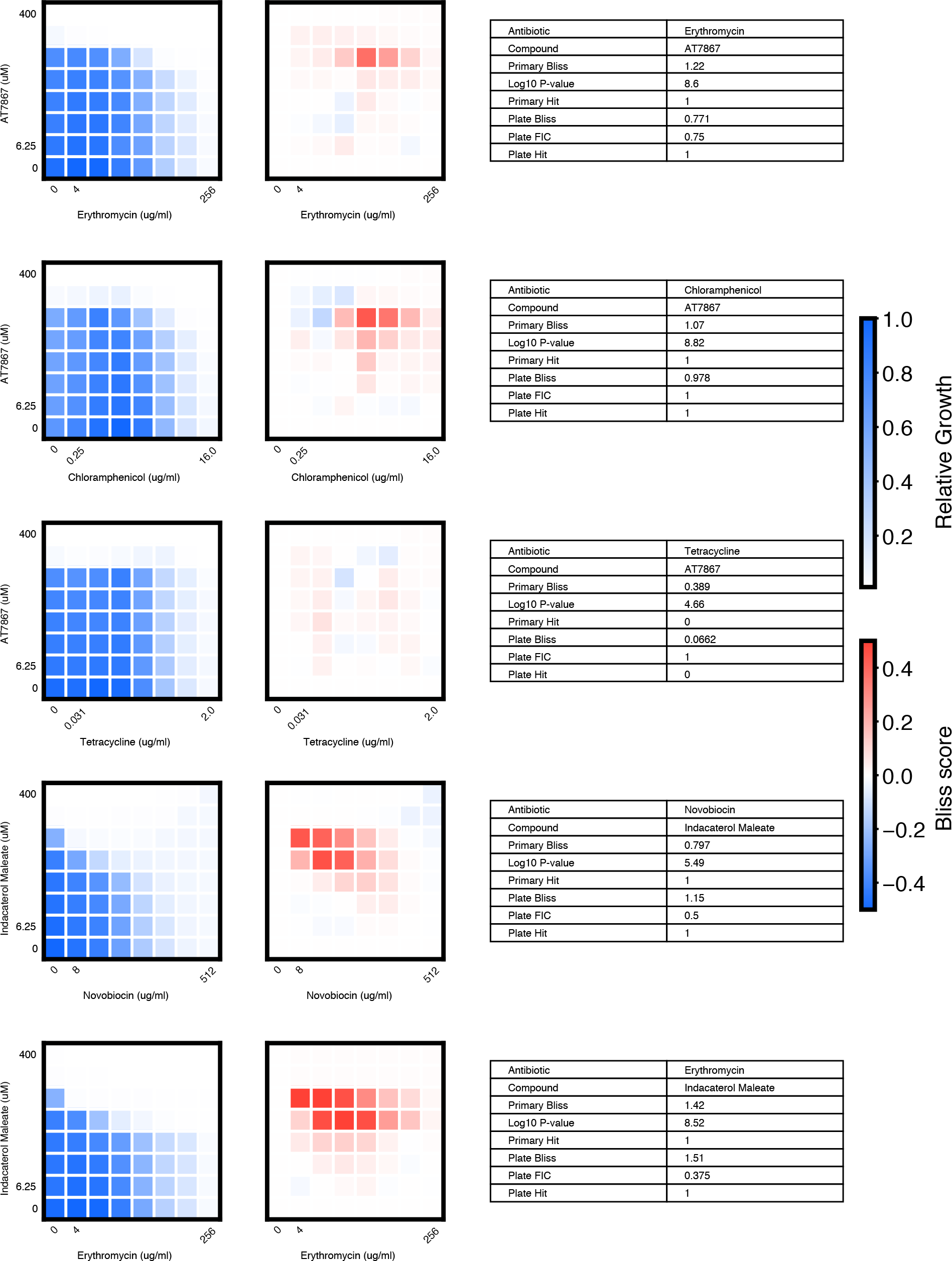
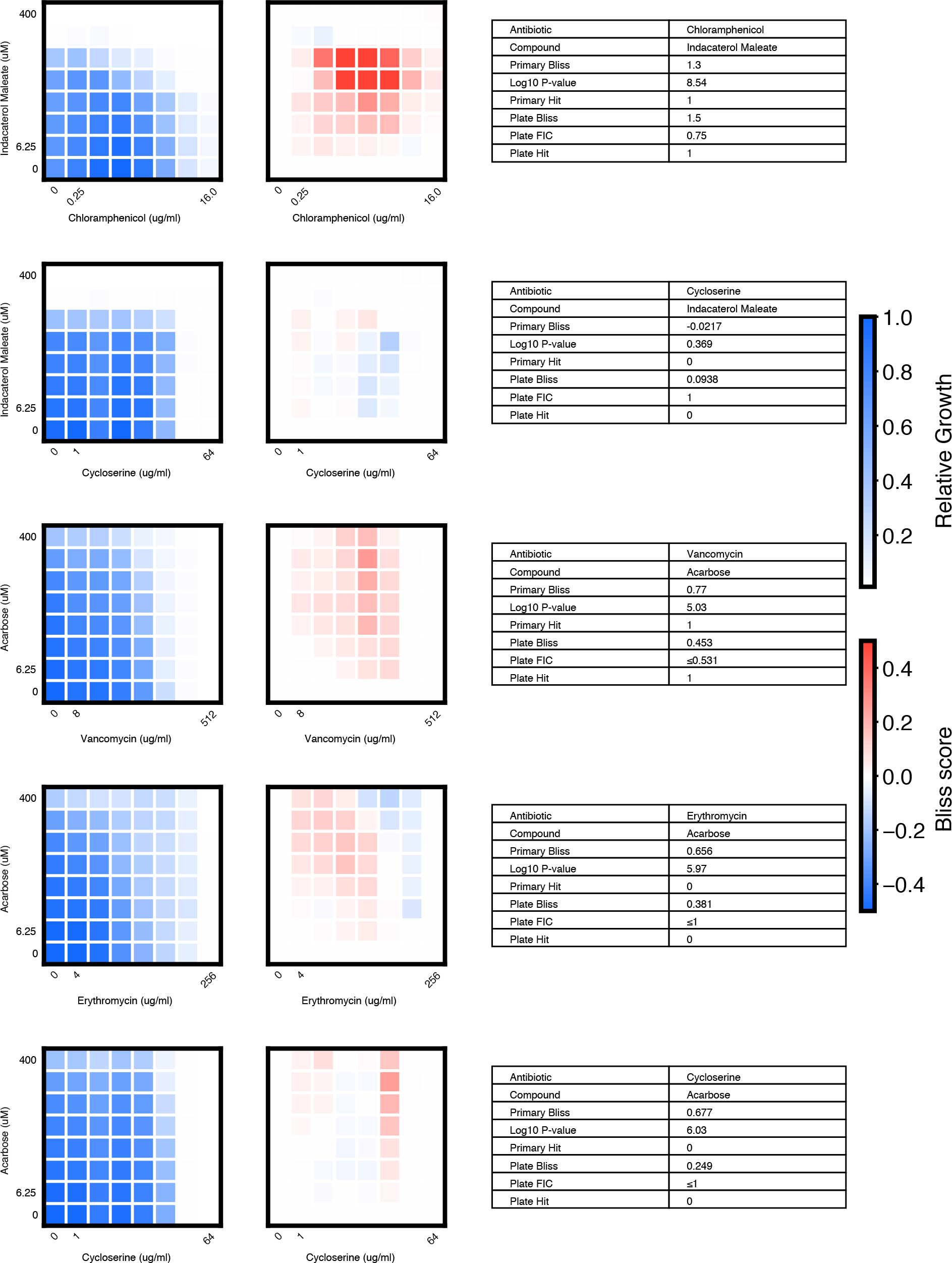
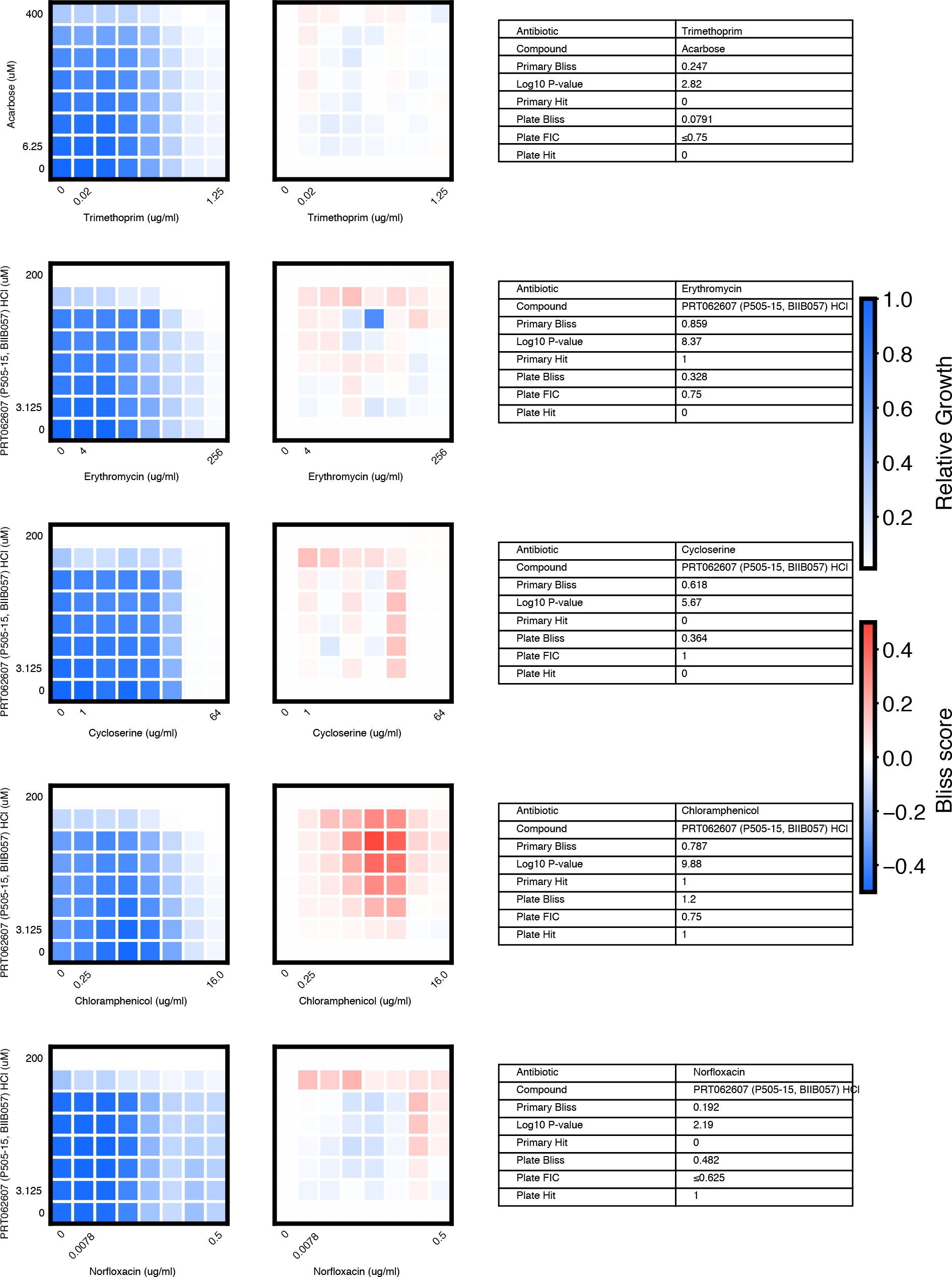
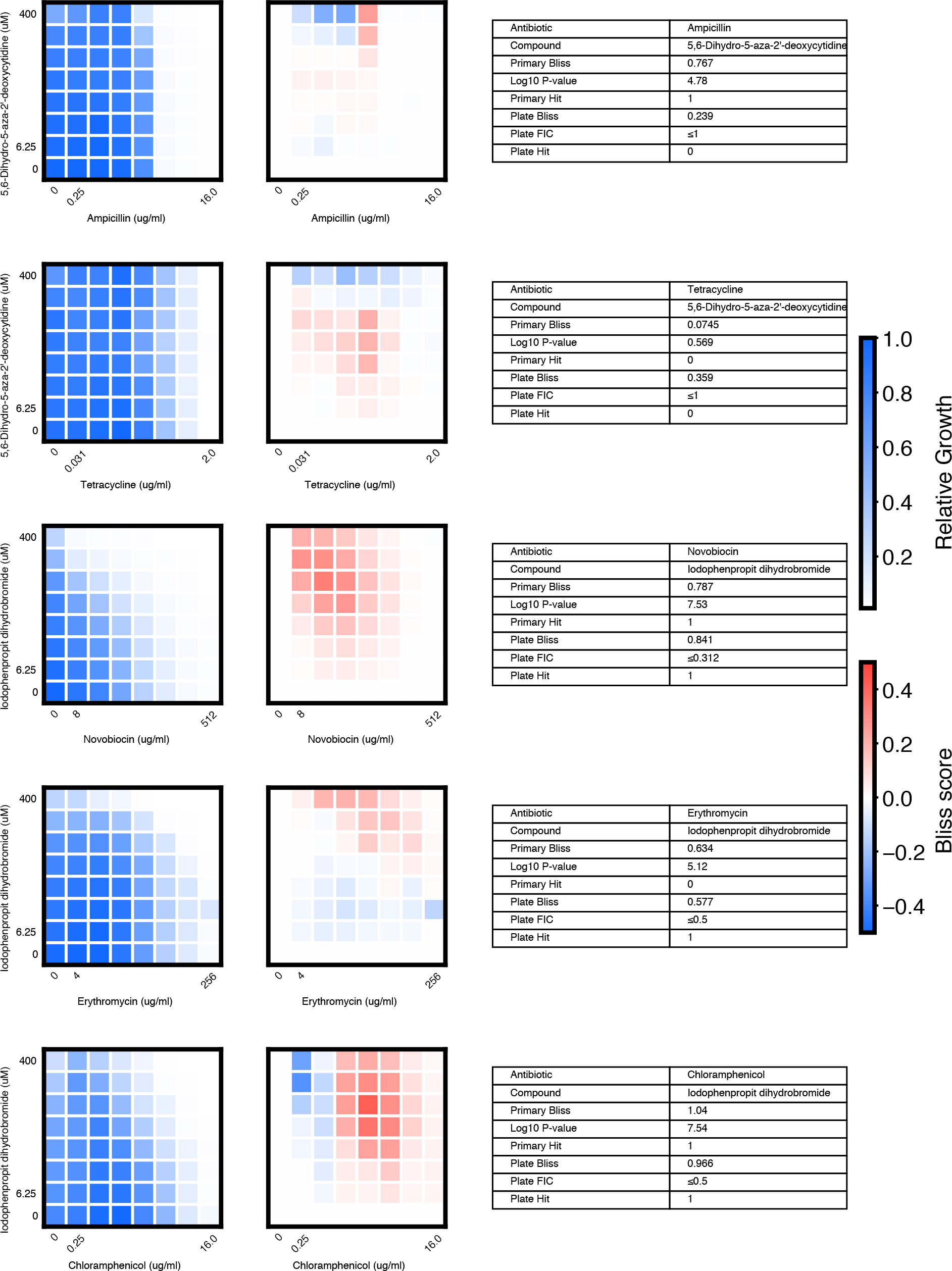
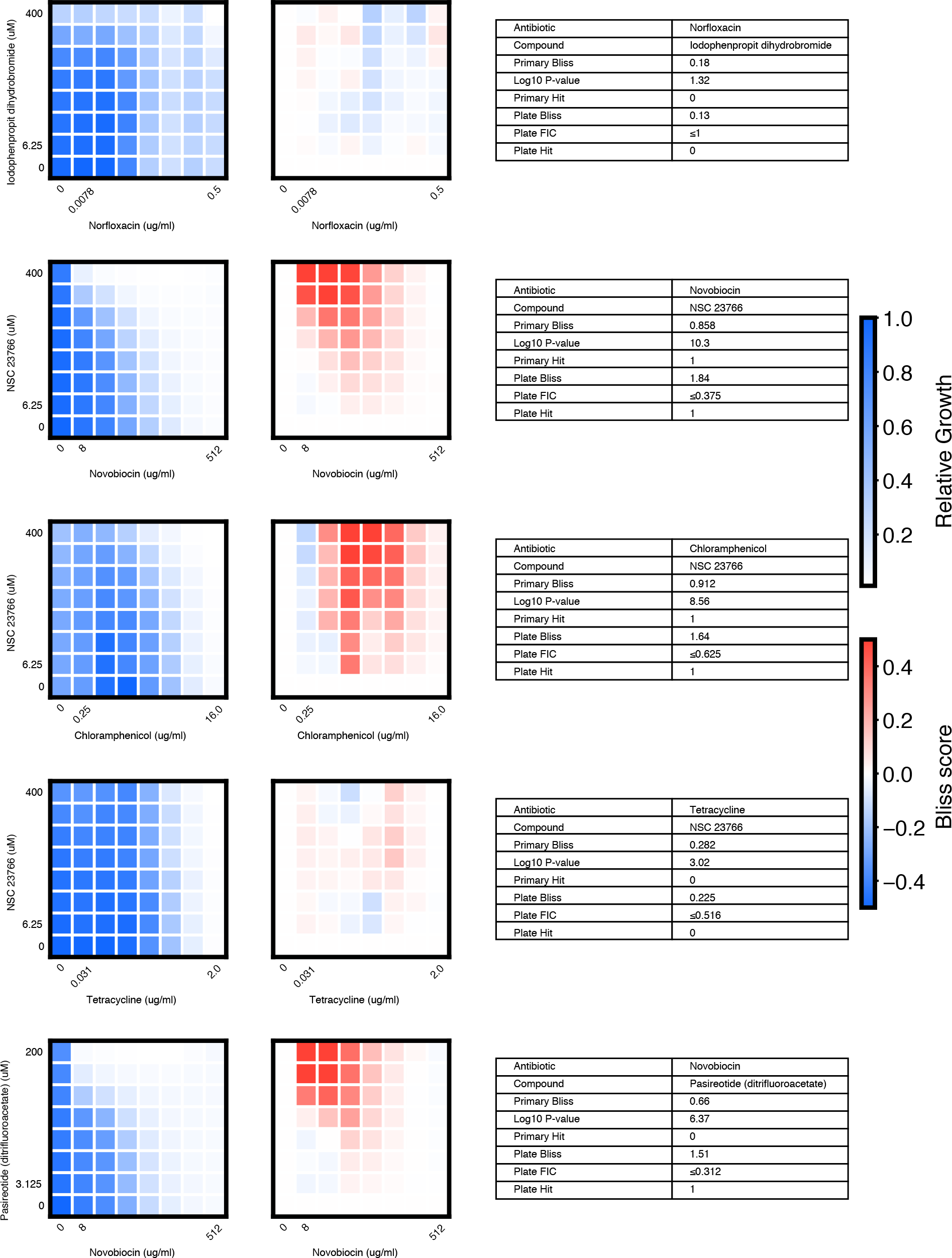
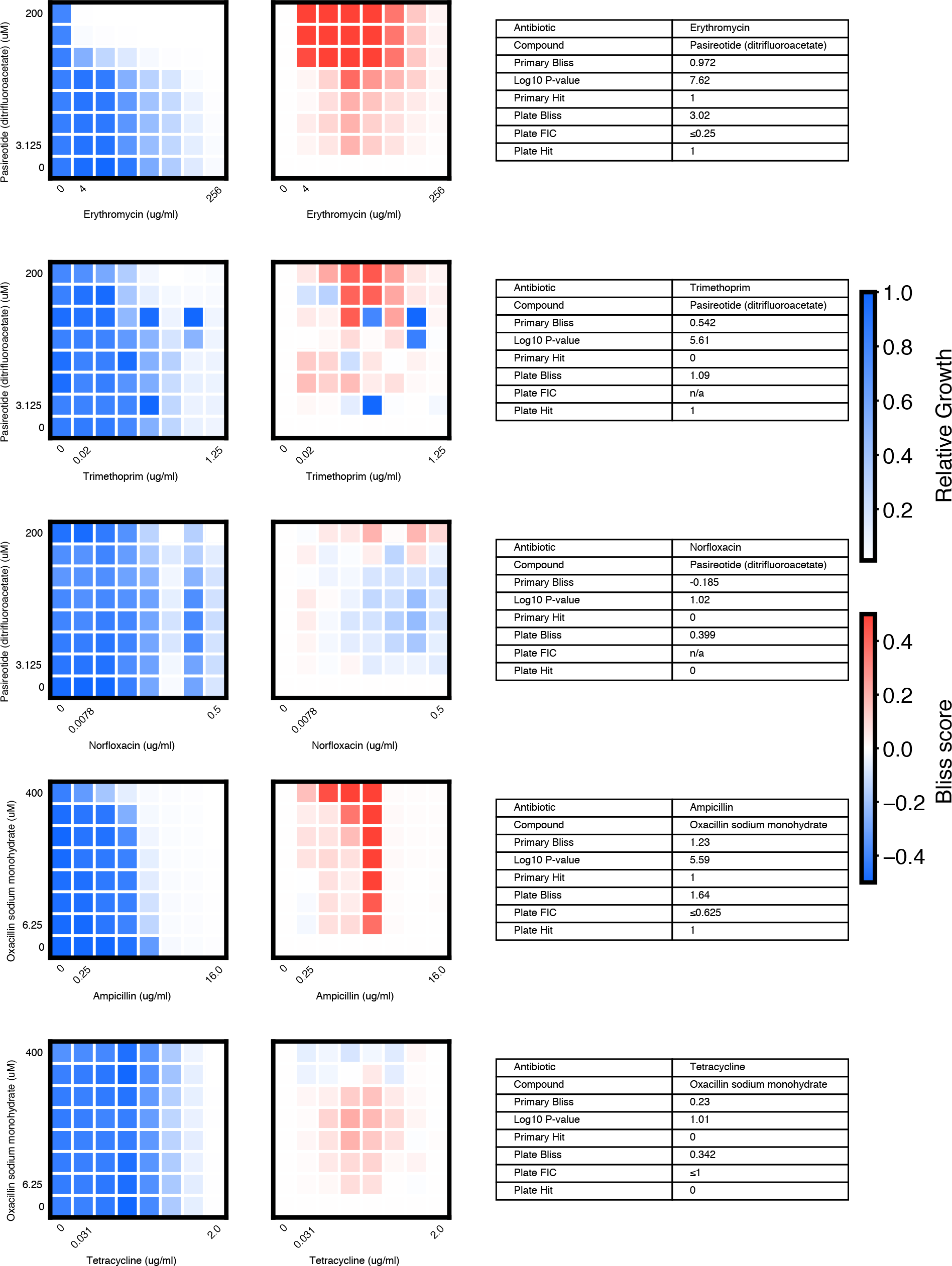
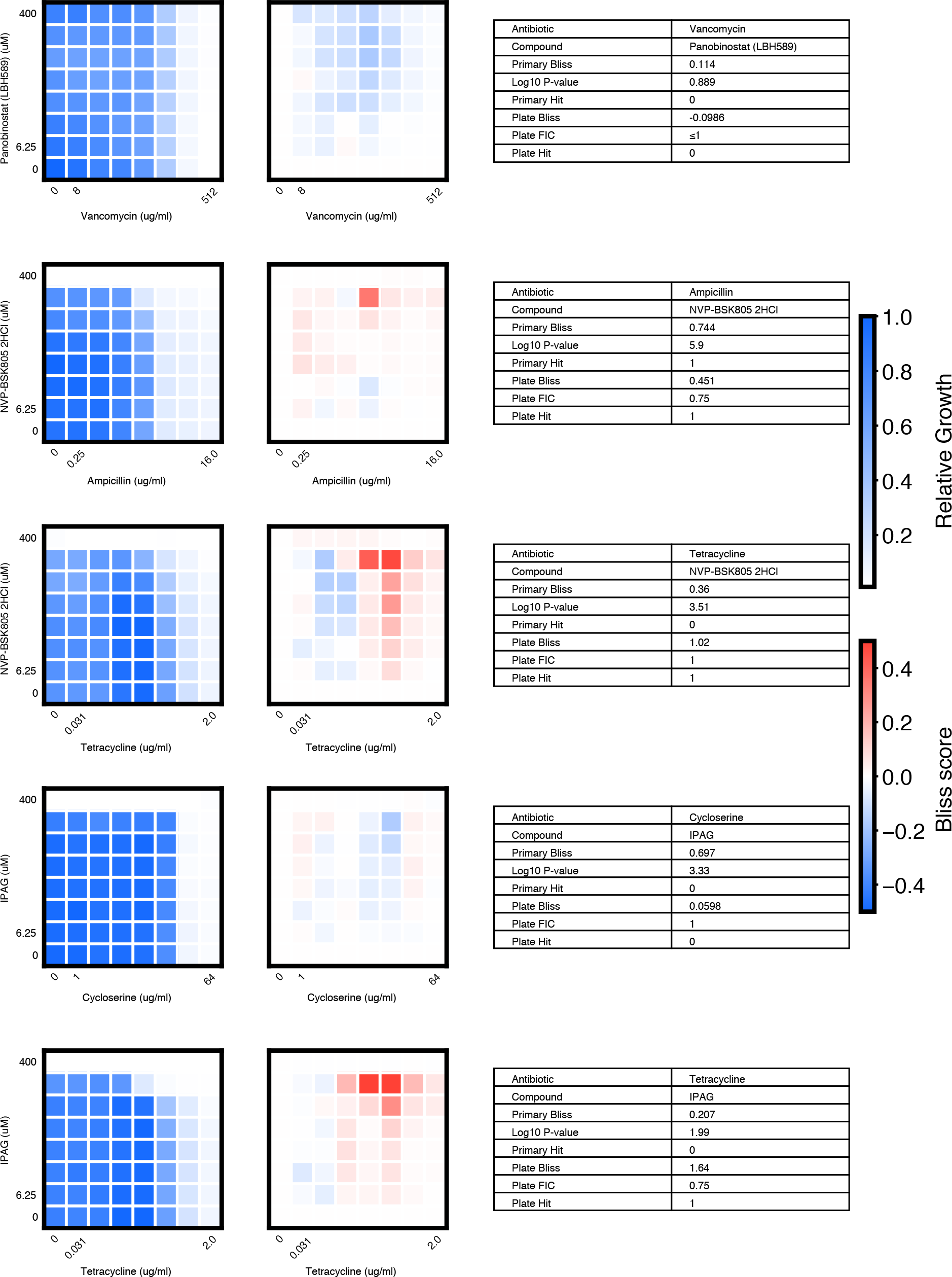
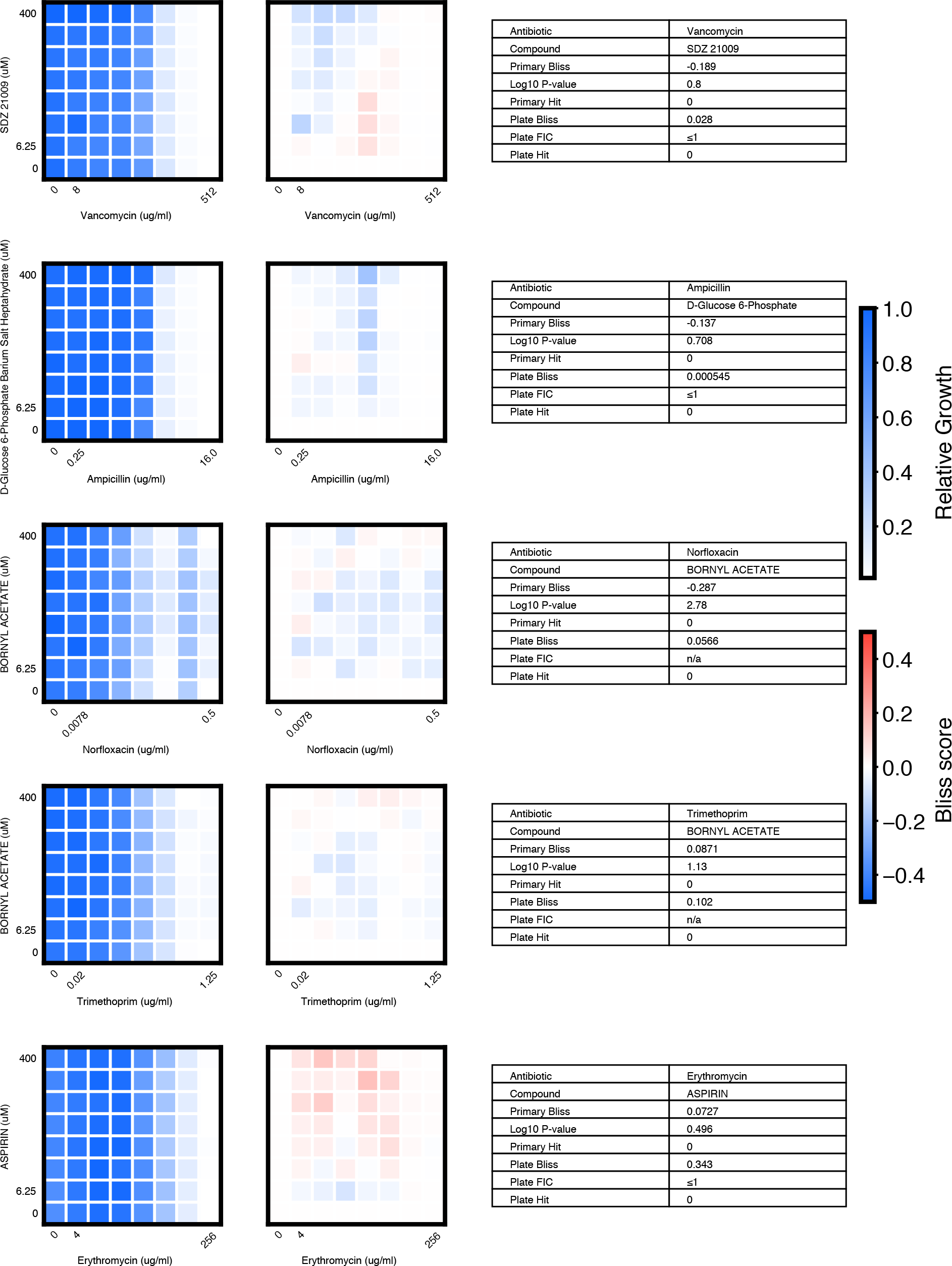
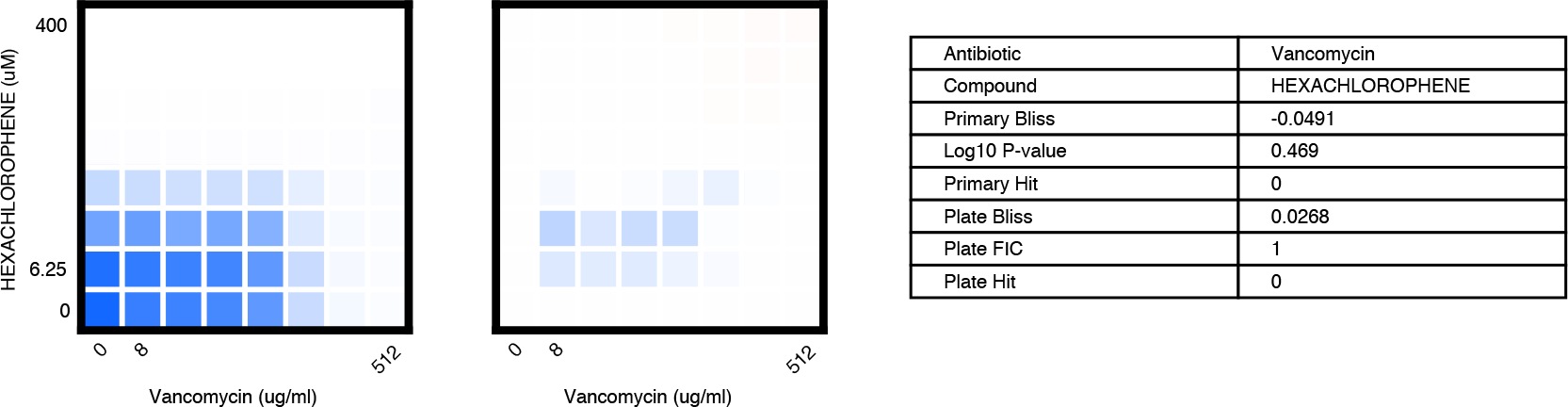
Checkerboard validation assays (full-scale screening phase). We chose 46 compound × antibiotic pairs to test the predictions made based on the primary screening data. For each checkerboard, the relative growth values (left panel), calculated Bliss Scores for each well position (middle panel), and a table summarizing the primary screening data and checkerboard synergy scores (Bliss Score and FIC) are shown (**materials and methods**).

**Figure S13.**
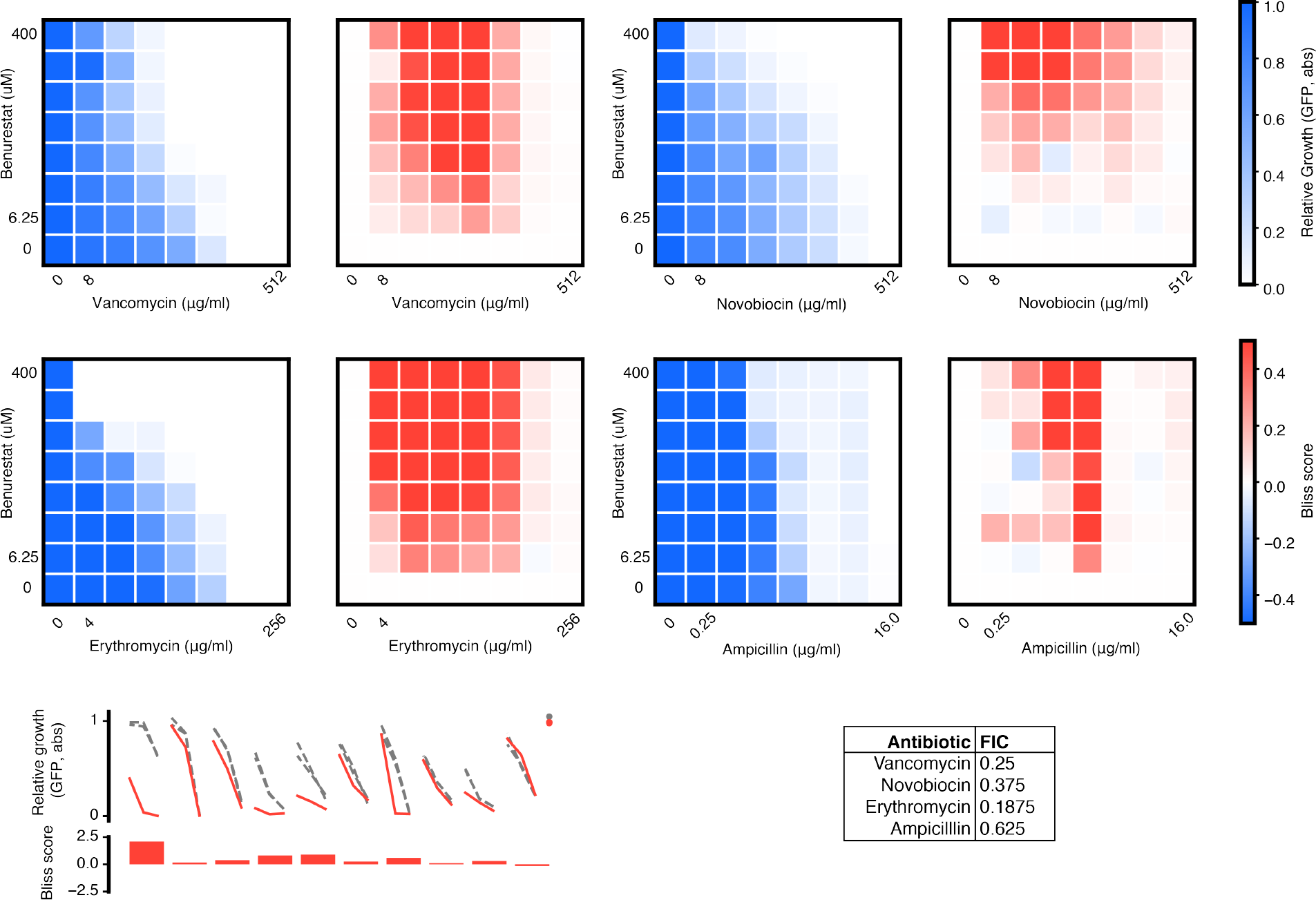
Checkerboard validation assays – Benurestat (pilot phase). We tested benurestat for synergies predicted with four antibiotics, measured by Bliss score and FIC (**materials and methods**). For each checkerboard, the relative growth values (left) and calculated Bliss Scores for each well position (right panel) are shown. Corresponding FIC values are shown in the table below. Primary screening data are also shown for comparison.

**Figure S14.**
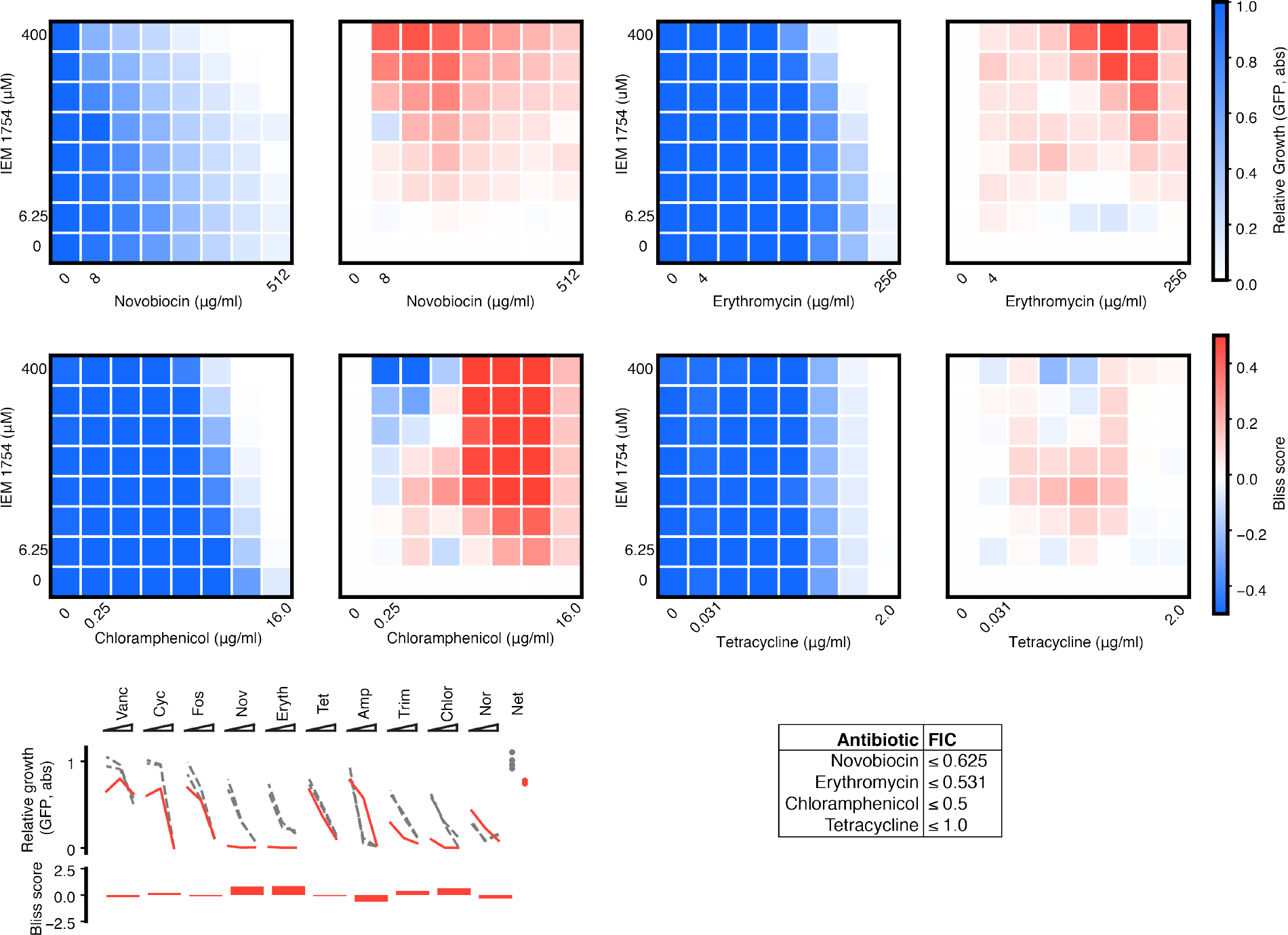
Checkerboard validation assays – IEM 1754 (pilot phase). We tested IEM 1754 for synergies predicted with three antibiotics, and one antibiotic predicted to show independent effects (tetracycline), measured by Bliss score and FIC (**materials and methods**). For each checkerboard, the relative growth values (left) and calculated Bliss Scores for each well position (right) are shown. Corresponding FIC values are shown in the table below. Primary screening data are also shown for comparison.

**Figure S15.**
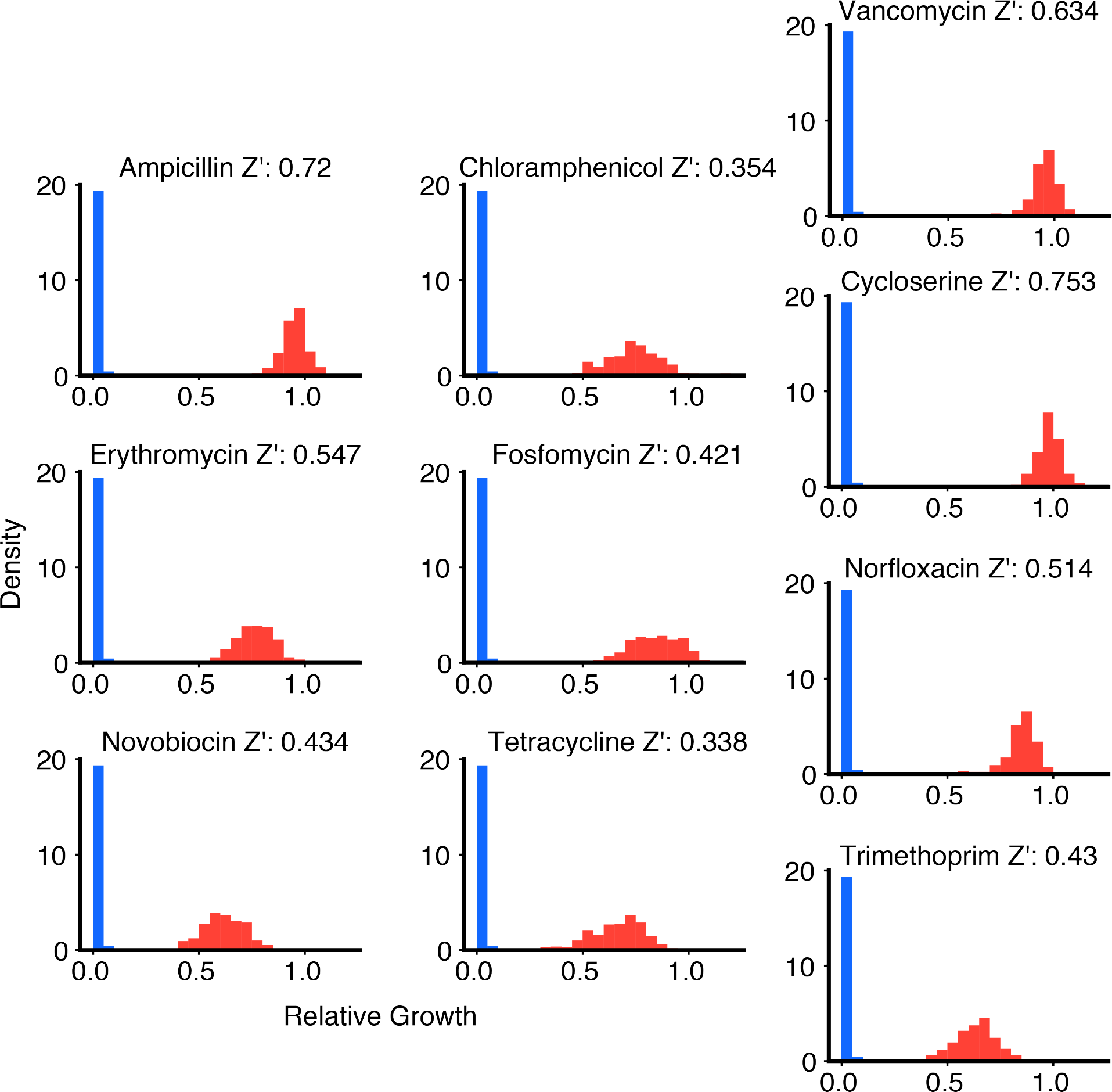
Screening assay performance for each antibiotic in panel. For each antibiotic in our panel, the dynamic range of the potentiation assay is the difference between the high relative growth values for the antibiotic at the lowest concentration tested (red histogram) and the low relative growth values observed under strong inhibition (cycloserine 16 μg/mL, blue histogram, same in all plots) (**materials and methods**) across all 108 chips analyzed. The difference between these distributions constitutes the antibiotic-specific dynamic range within which potentiation can be detected. To quantify assay performance, we computed the Z-factor (Z’, displayed in title of each plot) between these two distributions for each antibiotic.

## Supplementary Movies

**Movie S1:** Pool of 1-nanoliter droplets (food coloring) loaded into a large-format (**Fig. S1**) microwell array using a P1000 micropipette.

**Movie S2:** Demonstration of microwell array chip loading and washing procedure on large-format array (**Fig. S1**).

**Movie S3:** Demonstration of droplets merging on the microwell array chip upon exposure to an AC electric field (**materials and methods**). The width of each microwell is 148.6 μm and the length is 271.4 μm (**Fig. S1**).

## Acknowledgements

Relevant data reported in this paper are attached in the supplementary figures and tables. The drug repurposing library screened here is extensively annotated online at clue.io/repurposing. The authors thank Navpreet Ranu and David Feldman for early discussions about microfluidic and encoding strategies. The authors also thank Deepan Thiruppathy and Jameson Kief for assistance; Stewart Fisher, Jonathan Stokes, Jason Yang, and Wesley Chen for discussions; the Hung Lab (Broad Institute) for bacterial samples and discussions; Chris Emig and Tommy Moriarty for assistance designing and fabricating custom pressure manifolds; Josh Bittker, Samuel Figueroa-Lazú, Anita Vrcic, and the Broad Institute Compound Management team for compound library formatting and quality control; Scott Sassone for production of videos; and the jupyter, numpy, scipy, scikit-image, scikit-learn, and pandas open source development teams. This work was supported in part by the National Science Foundation Graduate Research Fellowship Program (A.K., J.K.), the MIT Institute for Medical Engineering and Science Broshy Fellowship (A.K.), a Career Award at the Scientific Interface from the Burroughs Welcome Fund (P.C.B.), an MIT Deshpande Center Innovation Grant, a Scialog seed grant from the Gordon and Betty Moore Foundation and the Research Corporation for Science advancement, and a Bridge Project grant from the Dana Farber/Harvard Cancer Center and the Koch Institute for Integrative Cancer Research at MIT. The Broad Institute and MIT may seek to commercialize aspects of this work, and related applications for intellectual property have been filed.

